# Preconfigured dynamics in the hippocampus are guided by embryonic birthdate and rate of neurogenesis

**DOI:** 10.1101/2022.05.07.491015

**Authors:** Roman Huszár, Yunchang Zhang, Heike Blockus, György Buzsáki

## Abstract

The incorporation of novel information into the hippocampal network is likely be constrained by its innate architecture and internally generated activity patterns. However, the origin, organization, and consequences of such patterns remain poorly understood. Here, we show that hippocampal network dynamics are affected by sequential neurogenesis. We birthdated CA1 pyramidal neurons with in-utero electroporation over 4 embryonic days encompassing the peak of hippocampal neurogenesis, and compared their functional features in freely moving, adult mice. Neurons of the same birthdate displayed distinct connectivity, coactivity across brain states, and assembly dynamics. Same birthdate hippocampal neurons were topographically organized, in that anatomically clustered (<500µm) neurons exhibited overlapping spatial representations. Overall, the wiring and functional features of CA1 pyramidal neurons reflected a combination of birthdate and the rate of neurogenesis. These observations demonstrate that sequential neurogenesis in embryonic development shapes the preconfigured forms of adult network dynamics.

## INTRODUCTION

The hippocampus plays a crucial role in the rapid encoding and storage of episodic memories.^1^ This function is thought to depend on its unique anatomical and functional organization.^2^ In contrast to the neocortex where receptive fields are topographically organized (i.e., physically nearby neurons have similar fields), hippocampal ‘place cells’ that represent the same or different parts of an environment are thought to be randomly distributed throughout the hippocampus.^3,4^ It has been suggested that the specific constellation of place cells representing a particular environment is established via activity-dependent plasticity,^5–7^ which allows the expression of place fields at arbitrary locations. In each novel environment, the hippocampus may call up an independent map or ‘chart’ to be associated with sensory inputs.^8^ The anatomical organization of the CA3 recurrent system is assumed to form a large random graph,^2,9–11^ endowing the hippocampus with a large storage capacity. Based on these anatomical and physiological observations, theoretical and computational models of hippocampal function have been developed using a framework consisting of randomly connected uniform principal neurons.^12–14^

However, several considerations point to the oversimplicity of the orthogonal chart organization based on random connectivity and, instead, indicate that hippocampal principal cells are organized into heterogeneous, parallel circuit modules, enhancing computational flexibility and supporting a rich repertoire of behaviors.^15,16^ In the narrow CA1 pyramidal layer, the sole cortico-fugal output of the hippocampus, important differences have been noted in the septo-temporal, medio-lateral and radial organization. Intrinsic physiological features, short- and long-range connectivity, and place field properties vary with anatomical position within the pyramidal layer.^17–20^ Recent work also indicates that this rich heterogeneity is coupled with a preservation of the individual properties of neurons and their assembly cooperation. For example, firing rates of pyramidal neurons are preserved across environments and brain states.^21^ Pyramidal neurons exhibit one to several place fields and follow an exponential distribution,^22^ a property that is preserved across environmental conditions and time.^23,24^ The emergence of place fields is biased towards locations with weak subthreshold drive,^25,26^ and is predictable from a preexisting correlation structure.^27–29^ These observations have led to the alternative view that structural organization in hippocampal networks gives rise to a reservoir of preconfigured activity patterns, which are available for matching with novel experiences.^27,29–32^ A potential source of these pre-existing states is embryonic development.^33^

To address the origin of the functional heterogeneity and preconfigured dynamics in the adult brain, we probed the functional consequences of developmental events that take place prior to behavioral experience. To examine how intrauterine development affects future hippocampal function, we birthdated CA1 pyramidal neurons with in-utero electroporation^34^ at four embryonic stages, and performed high density silicon probe recordings to compare their functional features in freely moving, adult mice. Same birthdate pyramidal neurons exhibited distinct connectivity, coactivity across brain states, and cell assembly dynamics. Spatial representations were topographically organized in same day born populations, in that anatomically clustered (<500µm) same birthdate neurons were functionally related. With the aid of a computational model, we show that preexisting correlations between same birthdate neurons interact with the rate of neurogenesis to shape the diversity of observed assembly patterns. Our findings demonstrate that sequential neurogenesis in embryonic development guides the preconfigured forms of adult hippocampal networks.

## RESULTS

### Identification and in-vivo recordings from birthdated CA1 pyramidal neurons

To label CA1 pyramidal neurons with distinct birthdates, we performed in-utero electroporation of ChR2-EYFP and tdTomato in mouse embryos at four prenatal stages (**Fig. 1A** and **Supplementary Fig. 1**). In adult brains, pyramidal neurons born on embryonic (E) days 13.5, 14.5, 15.5 and 16.5 occupied broadly overlapping, yet distinct sublayers spanning the deep-to-superficial axis of the pyramidal layer (**Fig. 1B-C**).^33^ To investigate the dynamics of birthdated pyramidal neurons in vivo, adult mice that underwent electroporation at different embryonic stages (E13.5, n=3; E14.5, n=3; E15.5, n=5; E16.5, n=3) were implanted with high density silicon probes and optic fibers targeting CA1. Pyramidal neurons were separated from interneurons based on waveform shape and bursting statistics (**Supplementary Fig. 2A,B**). Birthdated pyramidal neurons were identified optogenetically by reliable, short latency discharge following 1.5-3ms light pulses (**Fig. 1D** and **Supplementary Fig. 2**; Methods). Waveform shapes of light responsive pyramidal neurons were no different from those of nonresponsive neurons (**Supplementary Fig. 2I**). Furthermore, light evoked spikes were systematically most similar to spontaneous spikes of their assigned cluster (**Supplementary Fig. 2C-F**), validating our identification of birthdated pyramidal cells in vivo.

**Figure 1.**
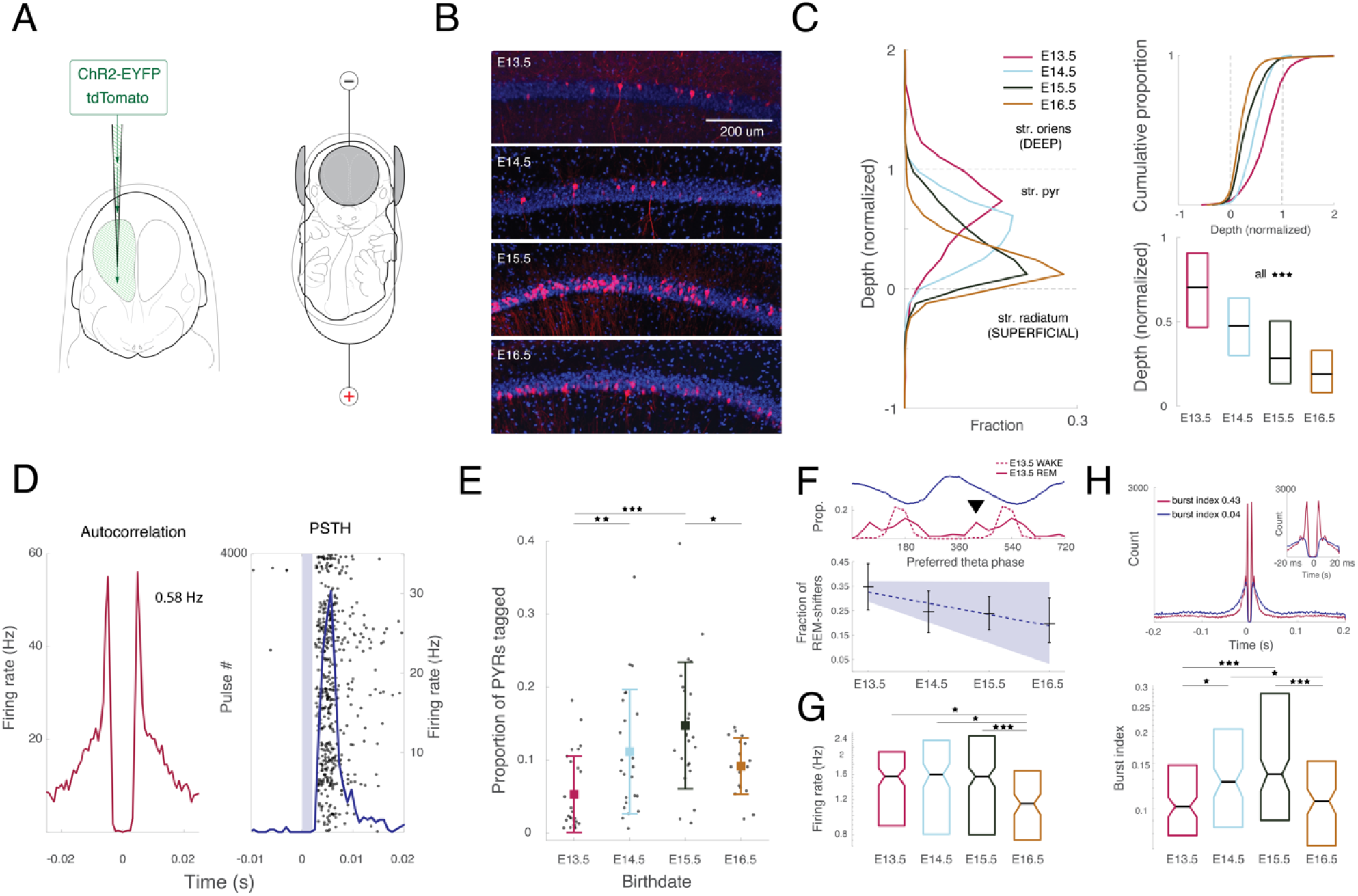
In vivo characterization of birthdated pyramidal neurons in mouse hippocampus. (**A**) Schematic of in-utero electroporation of DNA plasmids (ChR2-EYFP and tdTomato) for birthdating pyramidal neurons. A triple electrode technique was used to create an electric field that carried the plasmids towards progenitors of the CA1 subregion of the hippocampus. (**B**) Example histological images with tdTomato expression resulting from in-utero electroporation at four embryonic (E) dates: E13.5, E14.5, E15.5, and E16.5. (**C**) Left: Depth distributions of neurons with different birthdates within the pyramidal layer (Methods; **Supplementary Fig. 1B**). Right: Cumulative distributions and box plot summary of the depth distributions. E13.5, 0.71 (n = 2657 tdTomato+ puncta); E14.5, 0.48 (n = 1993); E15.5, 0.28 (n = 7749); E16.5, 0.19 (n = 9173). Median, Kruskal-Wallis: H=4.5e+03, df=3, p=0. Left: Spike autocorrelogram of a pyramidal neuron identified on the basis of waveform shape and bursting statistics. Right: Raster plot of the neuron’s optogenetic light (2ms, light blue) responses and superimposed peristimulus time histogram (PSTH, blue). (**E**) Proportion of recorded pyramidal cells tagged per session at different embryonic stages ^35^. Dots denote individual recording sessions. E13.5, 5.3±5.2% (n = 26); E14.5, 11.16±8.5% (n = 24); E15.5, 14.74±8.7% (n = 24); E16.5, 9.16±3.8% (n = 18). Mean±SD, ANOVA: F(3,88) = 7.82, p = 6e-04. (**F**) Top: Distribution of preferred theta phases of E13.5 pyramidal neurons (n=95) during waking and REM sleep, with an overlay showing the average LFP from two consecutive theta cycles. Arrowhead: REM sleep-induced theta phase shifting of spikes. Bottom: Fraction of theta-modulated birthdated pyramidal neurons that shifted their firing phase toward the peak of theta during REM sleep. E13.5, 34.7% (n = 95 neurons); E14.5, 24.5% (n = 106); E15.5, 23.8% (n = 143); E16, 19.7% (n = 76). Slope of regression line = -3.45, p=0.0374 (one tailed). Black crosses: mean±95% bootstrapped confidence interval (CI). Blue line and shading: linear regression±95% bootstrapped CI. (**G**) Firing rate distributions at different birthdates. E13.5, 1.57Hz (n=144); E14.5, 1.6Hz (n=132); E15.5, 1.57Hz (n=233); E16.5, 1.14Hz (n=115). Median, Kruskal-Wallis: H = 15.6604, df=3, p=3.6e-03. (**H**) Top: Autocorrelograms of two example pyramidal neurons with similar firing rates, but different burst propensities. Bottom: Burst indices (spike count at 2-5ms lags, divided by the spike count at 200-300ms lags) at different birthdates. E13.5, 0.101 (n=144); E14.5, 0.127 (n=132); E15.5, 0.136 (n=233); E16.5 = 0.107 (n=115). Median, Kruskal-Wallis: H=21.4329, df=3, p = 2e-04. *p<0.05, **p<0.01, ***p<0.001, n.s., not significant. For exact p-values of multiple comparisons, see **Supplementary Table 1**.

The fraction of light responsive pyramidal neurons at each birthdate was consistent with the previously observed bell-shaped wave of neurogenesis peaking between E14 and E15 (**Fig. 1E**).^35,36^ Earlier born pyramidal neurons had higher firing rates (**Fig. 1G**), and received the strongest drive from entorhinal cortex, as reflected indirectly by the largest fraction of pyramidal neurons whose theta phase preference shifted in REM sleep (**Fig. 1F**).^18,37,38^ In contrast, the propensity to fire in bursts displayed an inverted-U function of birthdate (**Fig. 1H**). This nonlinear relationship suggests a potential decoupling of the effect of anatomical positioning in the pyramidal layer from the effect of birthdate-related physiological parameters.

### Common birthdate shapes microcircuit arrangement and cofiring statistics of CA1 pyramidal neurons

In contrast to the neocortex, where clonally related sister pyramidal neurons synapse onto each other,^39^ CA1 exhibits negligible excitatory recurrence. To study the microcircuit arrangement of birthdated pyramidal neurons in our data, we focused on monosynaptic connectivity between pyramidal neurons and putative interneurons (**Fig. 2A**).^40^ Pyramidal neurons with the same birthdate (SBD) converged onto postsynaptic interneurons more strongly than neurons of different birthdates (DBD), and the magnitude of effective synaptic coupling (spike transmission probability) showed a nonlinear, bell-shaped relationship with birthdate (**Fig. 2B-E**).Furthermore, spike transmission probability peaked at presynaptic firing frequencies in the gamma range (15-23ms presynaptic interspike intervals) (**Fig. 2D**). Overall, SBD pyramidal cells projected onto common postsynaptic interneurons, which they drove most effectively at a timescale associated with assembly organization in the hippocampus.^41^

**Figure 2.**
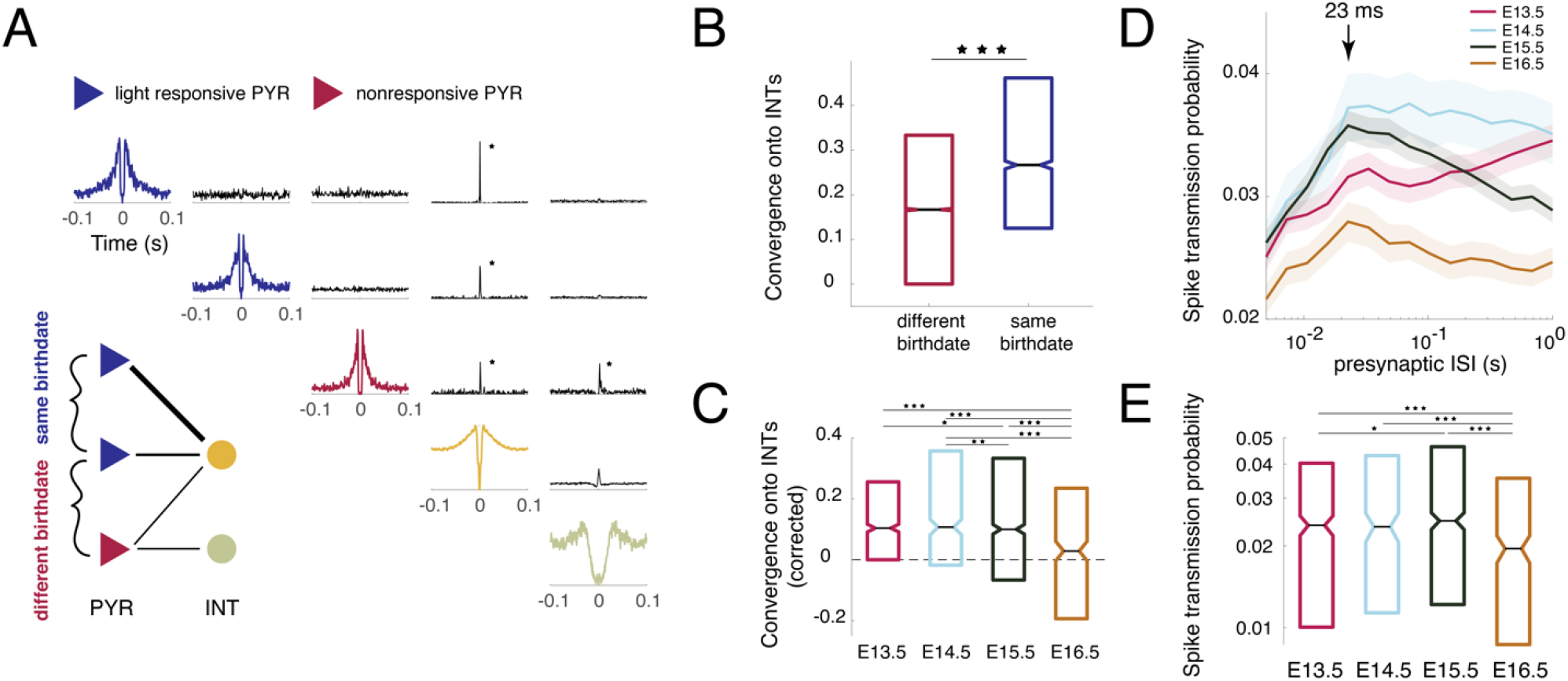
Microcircuit arrangement of birthdated pyramidal neurons. (**A**) Autocorrelograms of two same birthdate (SBD) pyramidal neurons (PYR; blue: light responsive), a different birthdate (DBD) neuron (red: non-responsive) and their convergence onto interneurons (INT). Significant peaks in the crosscorrelogram at 1-3 ms time lags revealed the presence of monosynaptic excitatory drive from pyramidal cells to interneurons.^40^ SBD pyramidal neurons converge onto the same postsynaptic interneuron. (**B**) Convergence onto shared interneurons of pairs of SBD neurons (blue, median=0.2667; n=3531), and of pairs of DBD neurons (red, median=0.1667; n=44201; p=4.9e-107; Wilcoxon rank-sum test). (**C**) Convergence of pyramidal cells onto interneurons at different birthdates. Values of SBD pairs at each birthdate were corrected by the median convergence of DBD pairs (dashed line). E13.5, 0.1039 (n=980 pairs); E14.5, 0.1071 (n=533); E15.5, 0.1 (n=1579); E16.5, 0.0285 (n=439). Medians, Kruskal-Wallis: H=71.2202, df=3, p=0. (**D**) Spike transmission (mean±S.E.M) onto postsynaptic interneurons at different presynaptic interspike intervals (ISI). Note short-term depression in spike transmission at presynaptic ISIs of <23 ms (i.e., cell assembly timescale).^41^ (**E**) Spike transmission probability at 15-23ms presynaptic ISIs, grouped by the birthdate of presynaptic pyramidal neurons. E13.5, 0.0238 (n=570); E14.5, 0.0235 (n=295); E15.5, 0.0247 (n=925); E16.5, 0.0195 (n=349). Medians, Kruskal-Wallis: H=16.1551, df=3, p=1.8e-03. *p<0.05, **p<0.01, ***p<0.001. For exact p-values of multiple comparisons, see **Supplementary Table 2**.

The microcircuit embedding of SBD neurons led us to explore their cofiring statistics. Crosscorrelograms pointed to greater synchrony between SBD than DBD pyramidal neurons (**Fig. 3A**), an observation that was appreciable in 10/12 animals (**Fig. 3B**). Furthermore, the time window of increased synchrony appeared to correspond with the average duration of sharp wave-ripples (SPW-Rs) (**Fig. 3B**), a brain state that engages large fractions of pyramidal neurons to fire.^42^ To quantify these observations, we focused on the firing statistics of pyramidal neurons during SPW-Rs (**Fig. 3C**). Both (SPW-R)-related firing rates and participation probability exhibited an inverted-U shaped relationship with birthdate (**Fig. 3D-E**). Consistent with the crosscorrelogram analyses, SBD pyramidal neurons exhibited greater pairwise correlations in SPW-Rs than DBD pyramidal neurons (**Fig. 3F**). This result held in 10/12 animals (**Supplementary Fig. 3**), and depended on neither firing rate nor cluster isolation quality differences (**Supplementary Fig. 4-6**).

**Figure 3.**
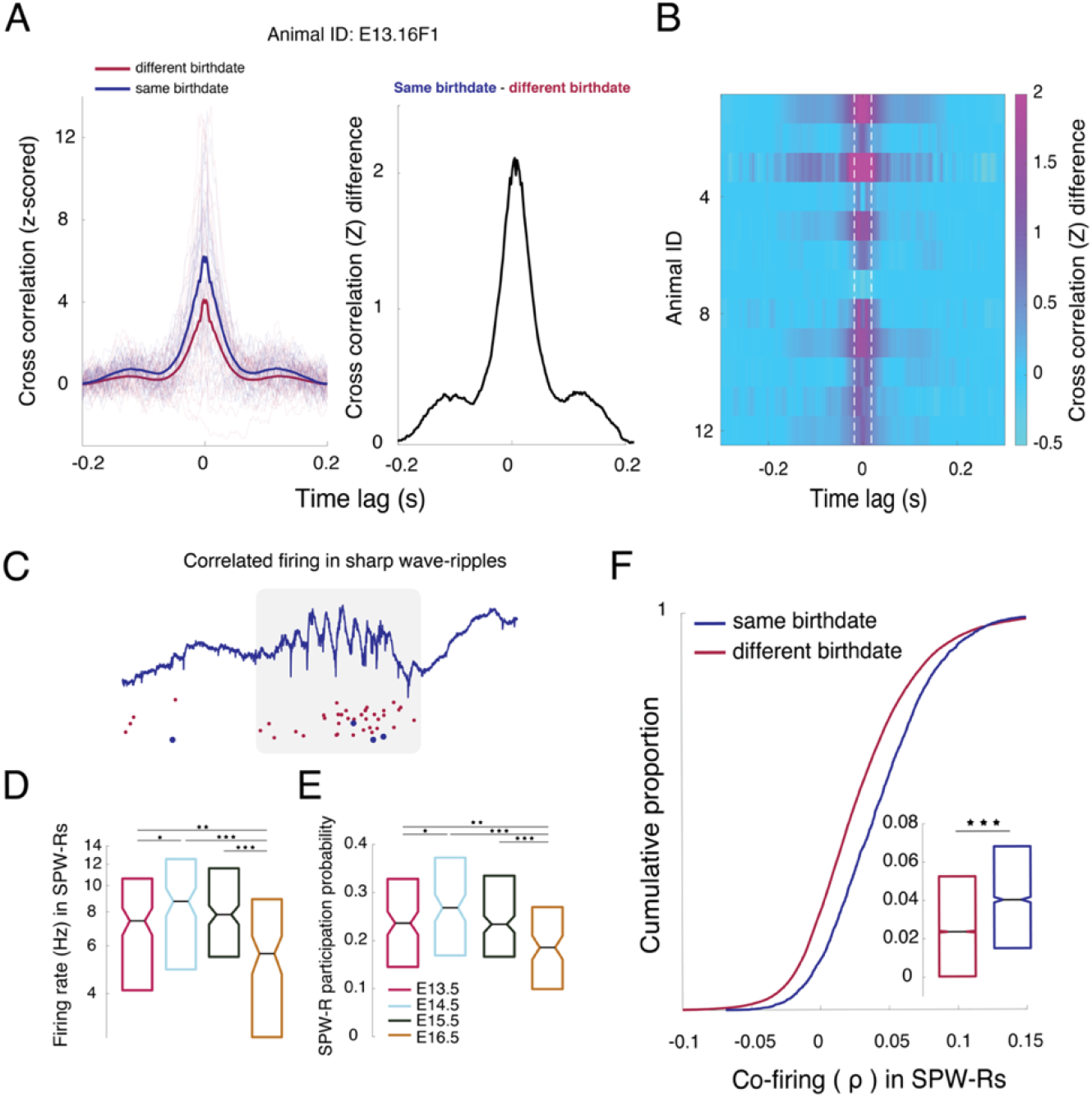
Sharp wave-ripple (SPW-R)-related cofiring of same birthdate pyramidal neurons. (**A**) Left: Z-scored crosscorrelograms for pairs of SBD and DBD pyramidal neurons recorded in an example animal (electroporated at E13.5). Bold lines are averages across all pairs (n = 977 and n = 13198, respectively), and thin lines are 10 example pairs from each group. Right: The difference between the average z-scored crosscorrelograms points to larger synchrony in the SBD group. (**B**) Difference between average z-scored crosscorrelograms between SBD and DBD pairs for each animal (row). Two E13.5 animals were excluded due to a lack of SBD pairs. White dotted lines indicate the median duration of sharp wave-ripples (SPW-Rs) across all recordings (36.4ms). (**C**) Example LFP and raster showing co-firing of SBD (blue dots) and other pyramidal neurons during a hippocampal SPW-R (grey rectangle). (**D**) Firing rates during SPW-Rs at different birthdates. E13.5, 7.42Hz (n=144); E14.5, 8.75Hz (n=132); E15.5, 7.82Hz (n=233); E16.5, 5.62Hz (n=115). Medians, Kruskal-Wallis: H=29.12, df=3, p=0. Fraction of SPW-Rs with at least one spike from a light-responsive neuron, shown across different birthdates. E13.5, 0.2363 (n=144); E14.5, 0.2684 (n=132); E15.5, 0.2341 (n=233); E16.5, 0.1856 (n=115). Medians, Kruskal-Wallis: H=26.1802, df=3, p=2e-04. (**F**) Pairwise spike count correlation in SPW-Rs for SBD (blue, median=0.0402; n=3531) and DBD pairs (red, median=0.0236; n=44201; p=1.4010e-105; Wilcoxon rank-sum test). *p<0.05, **p<0.01, ***p<0.001. For exact p-values of multiple comparisons, see **Supplementary Table 2**.

### Birthdated pyramidal neurons join assemblies with distinct dynamics

Given their tendency to converge on shared postsynaptic interneurons and to cofire during SPW-Rs, we hypothesized that SBD pyramidal neurons form functional microcircuits, reflected by distinct assembly dynamics. To study cell assemblies, we performed independent component analysis (ICA) on the Z-scored spike matrix of pyramidal neurons to extract patterns of higher order cofiring (**Fig. 4A-B**).^43^ Pyramidal neurons with large independent component (IC) weights (>2 SD) were considered assembly members, and individual assemblies were grouped by the birthdate of their members (Methods). Pairs of assembly members cofired in SPW-Rs more prominently than pairs of assembly non-members (**Fig. 4C**).^44^ Interestingly, assembly member cofiring with assembly non-members depended nonlinearly on birthdate; specifically, (SPW-R)-related firing patterns of pyramidal neurons that were members of assemblies associated with earliest (E13.5) and latest (E16.5) birthdates were the most segregated (i.e., lowest correlations) from the firing patterns of assembly non-members (**Fig. 4C**). To further explore the heterogeneity of assembly activity, we obtained time-resolved estimates of assembly expression by projecting each IC onto the Z-scored spike matrix (**Fig. 4D**). Timestamps associated with significant peaks in the resulting timeseries were considered moments of assembly expression and analyzed further (Methods). Assemblies were expressed in SPW-Rs at rates that depended nonlinearly on assembly member birthdate; in particular, assemblies with members born at intermediate birthdates (E14.5) were expressed at higher rates than those with members born at early (E13.5) and late (E16.5) birthdates. This outcome agrees with our pairwise analysis (**Fig. 4C**) and suggests that the microcircuits which bias assembly membership of SBD neurons vary across birthdates.

**Figure 4.**
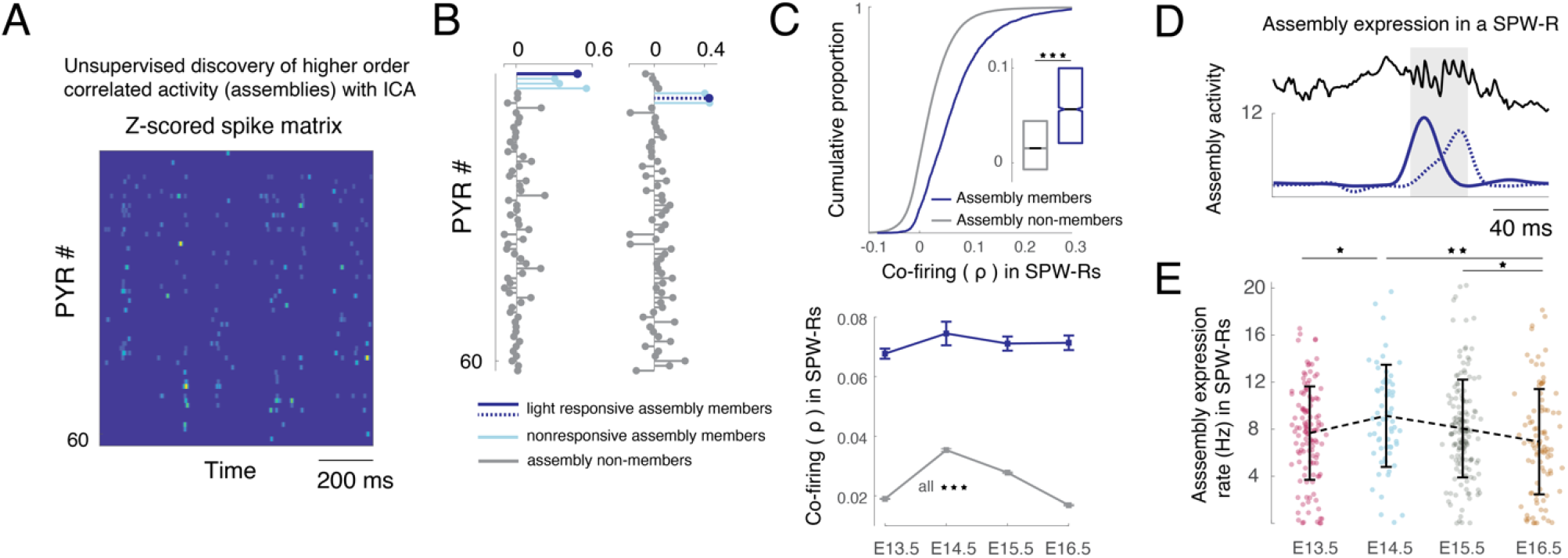
Distinct dynamics of cell assemblies with SBD neuron members. (**A**) Example z-scored spike matrix from 61 simultaneously recorded pyramidal neurons. Independent component analysis (ICA) was performed to identify assemblies, groups of neurons displaying prominent co-activation.^43^ (**B**) Example weights associated with two ICs. Pyramidal neurons with weights exceeding 2 standard deviations were considered assembly members. Blue, light responsive (birthdated) assembly members. Light blue, nonresponsive assembly members. Grey, assembly non-members. (**C**) Top: Pairwise spike count correlations in SPW-Rs between members of the same assembly (blue, median=0.0565; n=3516), and between assembly member and non-member neurons (grey, median=0.0154; n=155053; p=0; Wilcoxon rank-sum test). Bottom: Same as above, grouped by the birthdate of assembly members. Assembly member v. non-member cofiring (grey): E13.5, 0.019±1.7e-04 (n=66831); E14.5, 0.035±4.5e-04 (n=12368); E15.5, 0.028±2.25e-04 (n=37304); E16.5, 0.017±2.23e-04 (n=38550). Mean±S.E.M, ANOVA: F(3,155049)=865.7479, p=0. (**D**) Time-resolved assembly expression (colors identical to those of birthdated assembly members in (B)) during a SPW-R, resulting from the projection of ICs onto each column of the Z-scored spike matrix. Significant peaks were taken as time points of assembly expression, resulting in a time series. (**E**) Assembly expression rate in SPW-Rs for assemblies with birthdated assembly members. E13.5, 7.67±3.97 (n=123); E14.5, 9.14±4.35 (n=65); E15.5, 8.05±4.16 (n=149); E16.5, 6.93±4.45 (n=95). Mean±SD, ANOVA: F(3,428)=3.7228, p=0.0248. Dots denote individual assemblies. *p < 0.05, **p < 0.01, ***p < 0.001, error bars=S.E.M, unless noted otherwise. For exact p-values of multiple comparisons, see **Supplementary Table 3**.

### Correlations between SBD neurons interact with the bell-shaped rate of neurogenesis to produce assemblies with diverse expression rates

The temporal profile of assembly expression rates **(Fig. 4E**) qualitatively resembled the bell-shaped wave of pyramidal neurons labeled at different birthdates (**Fig. 1E**). A potential explanation of this similarity is that assembly dynamics and pairwise correlations are biased by intrinsic single cell differences in SPW-R related firing, which also exhibited a bell-shaped pattern with birthdate (**Fig. 3D-E**). An alternate explanation is that the rate of neurogenesis interacts with a correlation rule.^45^ To distinguish between these hypotheses, we constructed a phenomenological model that allowed firing rates and pairwise correlations to be tuned independently (**Fig. 5A-B** and **Supplementary Fig. 7A**; see Methods).^46^ The number of neurons was set to follow a bell-shaped function of birthdate (**Fig. 1E** and **Supplementary Fig. 7B**) and firing rates across birthdates were set according to the empirically observed distributions in SPW-Rs (**Fig. 3D** and **Supplementary Fig. 7C**). When pairwise correlations between SBD neurons were absent, the model failed to generate assembly expression rates comparable to data (**Fig. 5C-D**). This suggests that the bell-shaped pattern of firing rates alone is not sufficient. In contrast, as we increased the strength of correlations, the model generated assembly dynamics that yielded a good match to those observed in data (**Fig. 5C-D** and **Supplementary Fig. 7D-E**). Furthermore, reproducing the observed patterns required correlations between neurons to decay at some minimum time constant (∼2h) of the difference of their birthdates (**Fig. 5D**). These results suggest that correlated activity between SBD neurons and a bell-shaped neurogenesis curve suffice to produce the assembly dynamics observed in data and offer a plausible mechanism for generating a diverse repertoire of assembly patterns.

**Figure 5.**
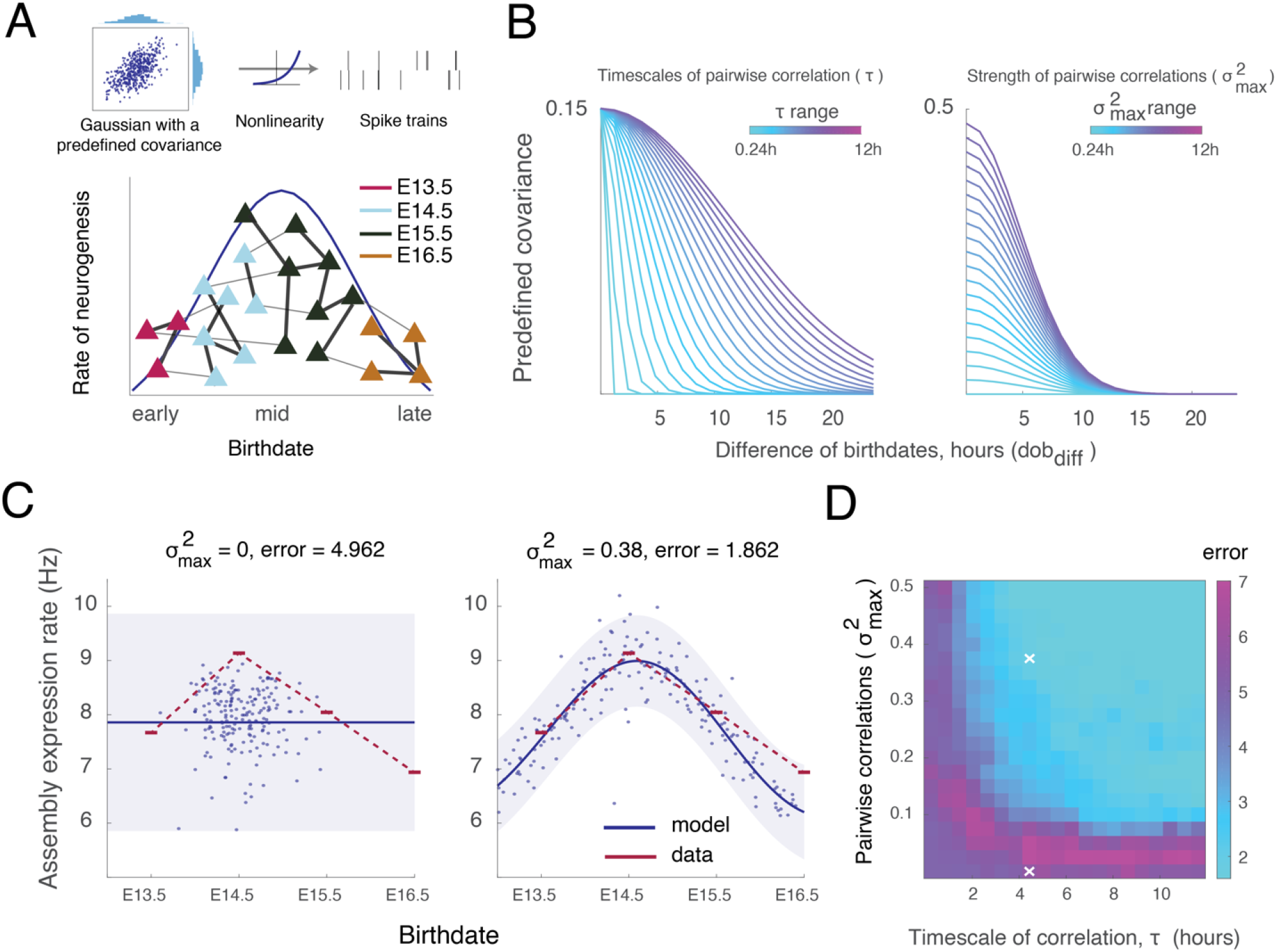
Correlations between SBD neurons interact with the rate of neurogenesis to produce the observed assembly dynamics. (**A**) Top: Schematic of a phenomenological model for exploring the statistical structure of assembly dynamics (**Supplementary Fig. 7**). Spikes were sampled from a transformed random process with independently tunable firing rates and correlations. Firing rates at different birthdates were set according to the firing rate distributions in SPW-Rs (**Fig. 3D**), and the strength of correlations between SBD pyramidal neurons was varied between simulations. Bottom: Schematic illustrating a hypothesized effect of neurogenesis rate and stronger functional correlations among SBD neurons. Large neurogenesis rates may give rise to more prominently expressed assemblies. Triangles, pyramidal neurons. Lines, strengths of pairwise correlations. (**B**) Illustration of the effect of free parameters τ and 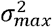 on the predefined covariance (Methods). Left: As τ increases, covariance decays more slowly as the difference of birthdates (*dob*_*diff*_) increases. Right: As 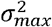 increases, the covariance decays from alarger initial value. In short, 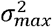 sets the strength of covariance between SBD neurons, whereas τ controls the rate at which it decays as the difference of birthdates increases, and implicitly defines the window during which neurons must be born in order to express preexisting correlations. (**C**) Simulated assembly expression rates at different birthdates (blue dots) yielded different qualitative fits to the observed assembly expression rates (red, average data as in **Fig. 4E**), depending on the strength of correlation 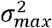 between SBD neurons. The timescale of correlation parameter τ was constant in the two examples. Goodness of fit was assessed as the negative log probability of the data under a nonlinear regression model summarizing the simulated assembly expression rates (blue line, posterior mean; shaded blue, posterior 95% confidence interval; Methods). (**D**) Negative log probability (error) matrix quantifying the model fit (as in (C)) as a function of the correlation strength between SBD neurons 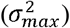, and the timescale of their correlation (τ). White crosses correspond to the examples in (C)).

### Same birthdate pyramidal neurons exhibited topographically organized spatial receptive fields

To explore the functional consequences of the developmentally set correlations, we trained adult mice (n = 8) that underwent in-utero electroporation to perform a place alternation task in a figure-8 maze with a 5s delay between choices (**Fig. 6A**). First, we observed that the strength of spike-theta phase coupling varied nonlinearly with birthdate. Specifically, intermediate birthdate pyramidal neurons (E14.5 and E15.5) locked to a broader range of theta phases (i.e., lower depth modulation) in the waking, but not REM state (**Fig. 6B**). Consistent with their cofiring during SPW-Rs (**Fig. 3F**), SBD pyramidal neurons exhibited greater correlations within theta cycles than DBD neurons (**Fig. 6C** and **Supplementary Fig. 4-6**). We therefore hypothesized that preexisting correlations of SBD neurons are a ubiquitous feature of the network and may constrain the expression of place-related firing.

**Figure 6.**
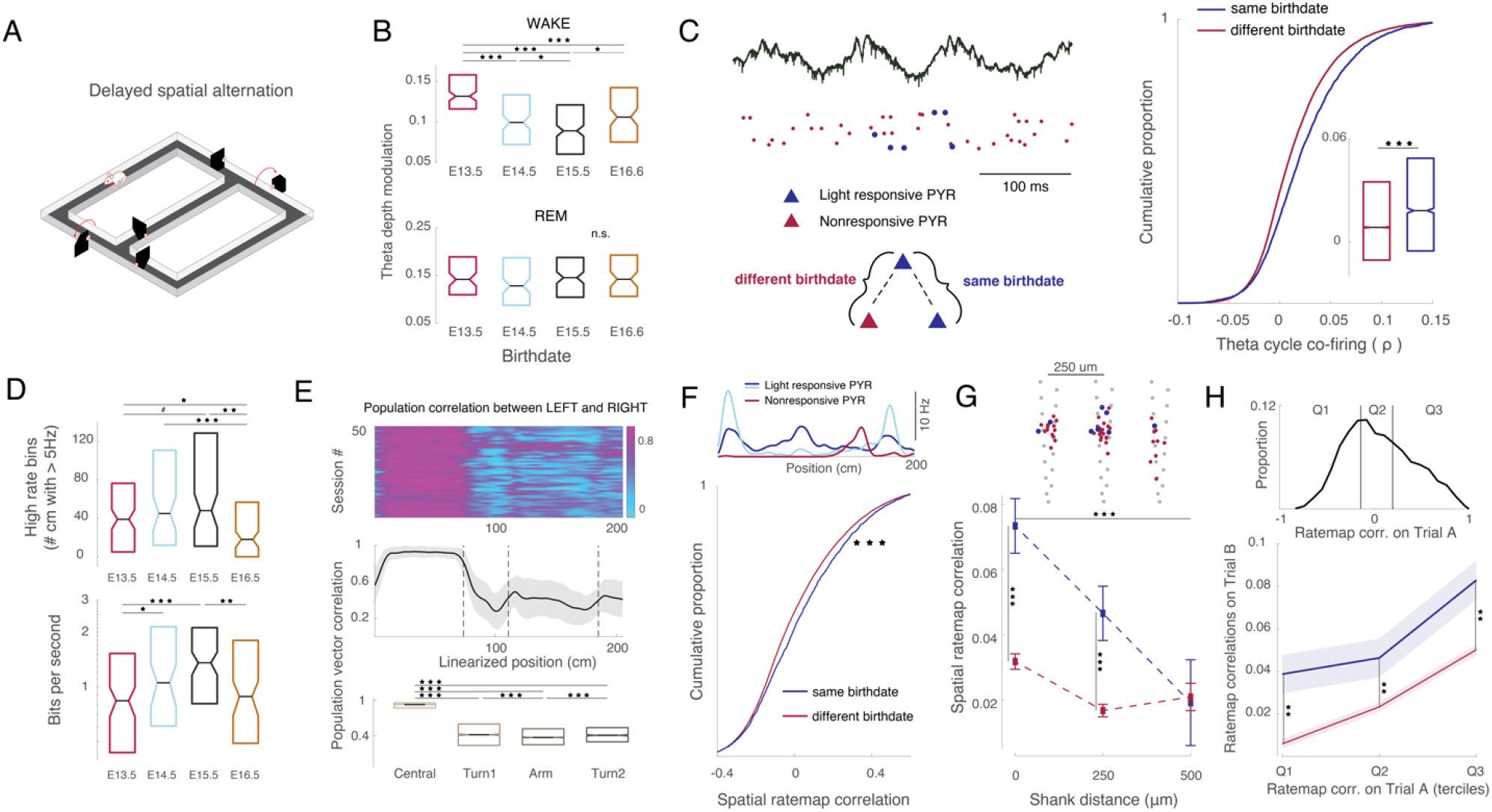
SBD neurons exhibit topographically organized spatial representations during behavior. (**A**) Schematic of a figure-8 maze. Water deprived animals alternated between left and right arms to obtain water reward. (**B**) Theta depth modulation of neurons at different birthdates. Top: During waking, E13.5, 0.12 (n=140); E14.5, 0.098 (n=127); E15.5, 0.09 (n=207); E16.5, 0.103 (n=101). Medians, Kruskal-Wallis: H=51.1573, df=3, p=0. Bottom: During REM, E13.5, 0.1443 (n=81); E14.5, 0.128 (n=99); E15.5, 0.1414 (n=118); E16.5, 0.1423 (n=73). Medians, Kruskal-Wallis: H=3.4993, df=3, p=0.4816. (**C**) Left: Example raster and LFP showing co-firing of SBD pyramidal neurons (blue) in the same theta cycle. Right: Pairwise spike count correlations in theta cycles between SBD (blue, median=0.0183; n=3531) and DBD pairs (red, median=0.0085; n=44201; p=4.47e-40; Wilcoxon rank-sum test). (**D**) Spatial coverage of firing during maze exploration at different birthdates. Top: Spatial extent (in cm) with high firing rate. E13.5, 39 (n=124); E14.5, 45 (n=76); E15.5, 48 (n=111); E16.5, 18 (n=115). Medians, Kruskal-Wallis: H=16.995, df=3, p=0.0022. Bottom: Spatial information score, in bits/s. E13.5, 0.84 (n=124); E14.5, 1.05 (n=76); E15.5, 1.35 (n=111); E16.5, 0.88 (n=115). Medians, Kruskal-Wallis: H=19.02, df=3, p=6e-04. (**E**) Population vector correlation between left and right trials as a function of the linearized position from the trial start. Significant decorrelation following the central arm (common to left and right trials) reflects distinct representations. Central, 0.93 (n=3922 population vector pairs); Turn1, 0.42 (n=1961); Arm, 0.375 (n=3922); Turn2, 0.411 (n=1060). Medians, Kruskal-Wallis: H=6713, df=3, p=0. (**F**) Spatial ratemap correlations between SBD (blue, median=0.017; n=2389) and DBD pairs (red, median=-0.0237, n=33705; p=2.9e-09; Wilcoxon rank-sum test). Top shows example spatial ratemaps for light responsive (blue) and nonresponsive (red) neurons. The central arm was excluded from each ratemap, and left and right ratemaps were concatenated for every pyramidal neuron. (**G**) Spatial ratemap correlation as a function of shank distance of recorded neurons. A two-way ANOVA revealed a main effect of birthdate (F(1,31544)=14.03, p=0.0002), shank distance (F(2,31544)=10.48, p=0), and their interaction (F(2, 31544)=3.4, p=0.0332). Top shows the positions of light-responsive (blue) and non-responsive (red) pyramidal neurons overlaid on the silicon probe recording sites (grey). Shank spacing was 250 µm. (**H**) Top: Ratemap correlation distribution between SBD pyramidal neurons, divided into terciles. Left and right trial ratemap correlations were calculated separately, and pooled. Bottom: Ratemap correlations in one trial type (left or right) as a function of the tercile in the other (right or left, respectively) for SBD and DBD pyramidal neurons. A two-way ANOVA revealed a main effect of birthdate (F(1, 68751)=32.07,p=0), tercile (F(2,68751)=25.52,p=0), but no interaction (F(2,68751)=0.39,p=0.68). #p<0.1, *p < 0.05, **p < 0.01, ***p < 0.001, n.s. not significant, all error bars = S.E.M. For all p-values of multiple comparisons, see **Supplementary Table 4**.

The spatial information content of spikes (measured in bits/s and as the number of spatial bins with high firing rate) was largest in pyramidal neurons with intermediate birthdates (E14.5 and E15.5; **Fig. 6D**).^19^ SBD neurons exhibited greater overlap in spatial tuning, as reflected in higher spatial ratemap correlations compared to DBD neurons (**Fig. 6F** and **Supplementary Fig. 4-6**). Surprisingly, the strength of spatial ratemap correlations depended on the anatomical distance between SBD neurons, in that spatial ratemaps of SBD pyramidal neurons ≥250μm apart exhibited systematically larger overlap than those of DBD neurons (**Fig. 6G** and **Supplementary Fig. 6**).

Changes in environmental context typically result in a global reorganization of spatial tuning. To explore this feature in birthdated populations, we compared their firing on the left and right arms of the figure-8 maze. Population firing rate vectors at linearized positions along the maze decorrelated between left and right trial types as soon as animals exited the common stem segment of the maze (**Fig. 6E**). Despite the different contexts across the two trial types, spatial ratemaps of SBD neurons tended to systematically reorganize closer together compared to those of DBD neurons (**Fig. 6H**). Overall, our results suggest that preexisting correlations between SBD neurons manifest in a topographically structured overlap of spatial representations, and this bias persists in the face of a change in environmental context.

### During behavior, birthdated pyramidal neurons join assemblies consisting of other SBD neurons

The above results are consistent with the possibility that during active behavior, birthdated pyramidal neurons are biased to fire in assemblies made up of other SBD pyramidal neurons. To detect assemblies of a held-out birthdated pyramidal cell, we identified its spikes during maze running and performed ICA on the remaining pyramidal neurons in the corresponding time bins (**Fig. 7A**). Assemblies that did not consist of other SBD pyramidal neurons were compared to those that did. We found that assemblies containing other SBD pyramidal neurons were more likely to be expressed surrounding the spikes of the held-out neurons (**Fig. 7B**), and their ratemaps overlapped more strongly with that of the held-out neuron (**Supplementary Fig. 7C-D**). Lastly, we compared the held-out pyramidal neuron’s theta cycle co-firing with SBD assembly members, DBD assembly members, and assembly non-members. Consistent with the (SPW-R)-related assembly results (**Fig. 4C**), held-out pyramidal neurons cofired with assembly members more strongly than with assembly non-members (**Fig. 7E**), and their cofiring with SBD assembly members was the strongest (**Fig. 7E**).

**Figure 7.**
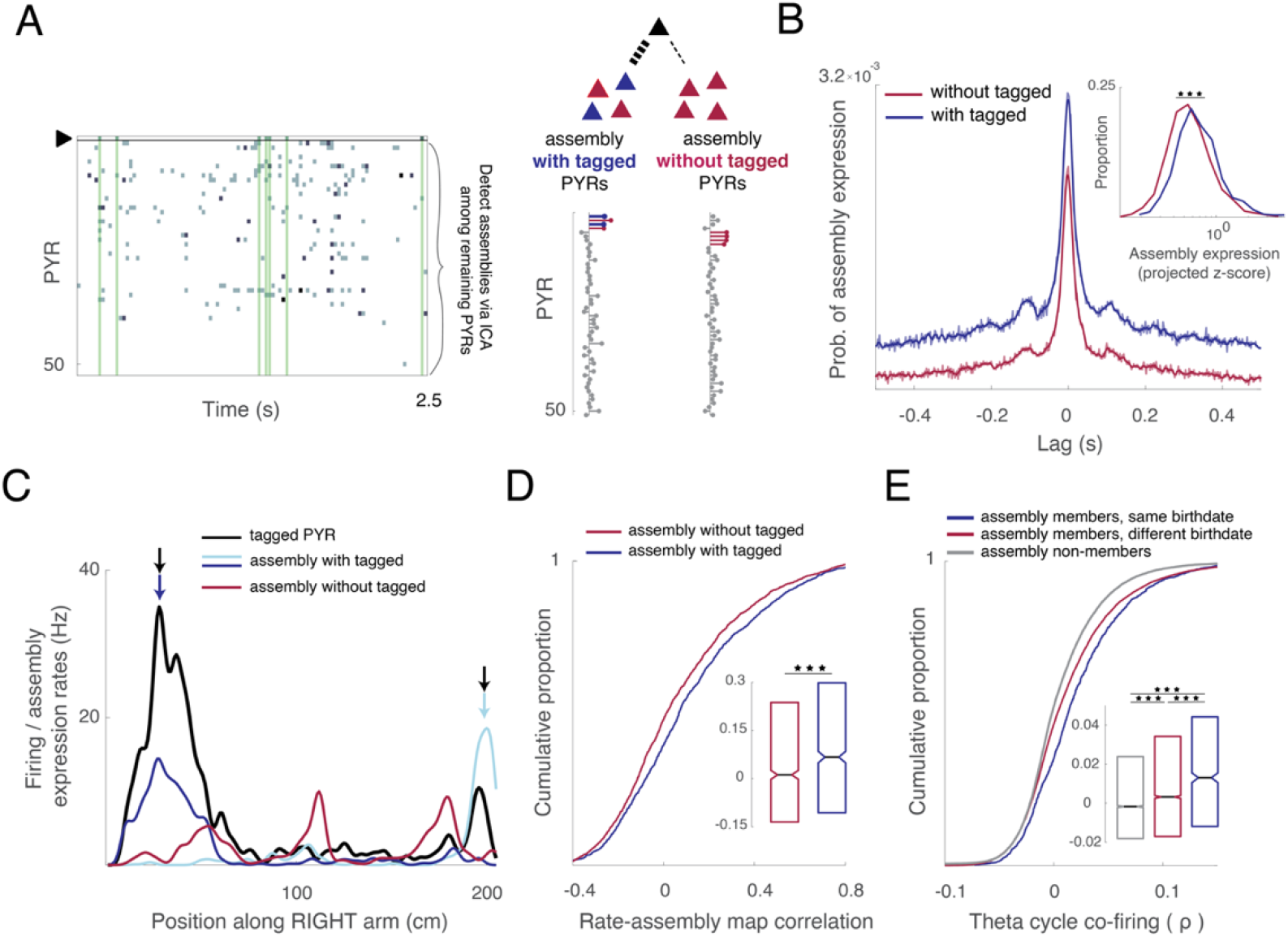
Birthdated pyramidal cells join assemblies consisting of other SBD neurons. (**A**) Left: 2.5s raster plot illustrating conditional assembly detection during behavior on the figure-8 maze (Methods). Assemblies were detected around the spikes of a held-out, birthdated pyramidal neuron (black triangle). ICA was performed on the remaining pyramidal neurons in 25ms time windows surrounding the spikes of the held-out neuron (vertical green bars). Right: Example independent components. Assembly members were determined as in **Fig. 4B**. Assemblies with other SBD neurons (blue) were compared to assemblies consisting exclusively of untagged neurons (red). (**B**) Probability of assembly expression at various time lags around the spikes of the held-out neuron. *Inset*, Distributions of assembly expression strengths at the time of the held-out neuron’s spike (blue, median=0.721, n=1259; red, median=0.61, n=1816; p=1.4e-23; Wilcoxon rank-sum test). (**C**) Firing ratemap of a held-out, birthdated pyramidal neuron (black), and assembly expression ratemaps for assemblies with tagged (dark and light blue) and untagged (red) members. Arrows indicate overlapping place fields. (**D**) Correlations between held-out neurons’ ratemaps and assembly expression ratemaps for assemblies with tagged (blue, median=0.067; n=1259) and untagged members (red, median=0.0116; n=1816; p=1.01e-05; Wilcoxon rank-sum test). (**E**) Pairwise spike count correlations in theta cycles between each held-out neuron and its SBD assembly members (blue), DBD assembly members (red), and assembly non-members (grey). w/ SBD members, 0.013 (n=1232); w/ DBD members, 0.0032 (n=12486); w/ non-members, -0.0017 (n=18417). Medians, Kruskal-Wallis: H=231.4156, df=2, p=0. *p < 0.05, **p < 0.01, ***p < 0.001, all error bars = S.E.M. For all p-values of multiple comparisons, see **Supplementary Table 4**.

## DISCUSSION

We show that sequential neurogenesis throughout embryonic development influences functional features of adult hippocampal activity patterns. Same birthdate pyramidal neurons exhibited overlapping projections onto postsynaptic interneurons, and strong correlations during sharp wave-ripples and theta waves. During behavior, preexisting correlations of same birthdate neurons exhibited overlapping place fields that depended on their anatomical proximity. Same birthdate neurons joined into cell assemblies whose frequency of expression scaled with the pool size of neurons that were born together. Our computational model suggests that pairwise correlations between same birthdate neurons interact with the bell-shaped wave of neurogenesis to generate diverse rates of assembly expression, as observed in our data. Altogether, we hypothesize that common developmental origin of hippocampal pyramidal cells biases their microcircuit arrangement, which both constrains and predicts single neuron and population firing features in the adult brain.

### Embryonic origin of functional organization in the CA1 pyramidal layer

The hippocampus can be conceived as a uniform large unit and its single-layer, archicortical organization is often contrasted to the modularly organized multi-layer neocortex.^2,47^ Yet, there appears to be a qualitatively similar Bauplan, with a gradual vertical expansion of pyramidal neurons from the hippocampus to subiculum to entorhinal cortex (paleocortex) and the six-layer isocortex.^48^ From this viewpoint, the CA1 pyramidal layer may be regarded as the ‘compressed’ version of the six-layer neocortex, displaying radial axis differences in input-output connectivity. This hypothesis resonates with the classic, two-sublayer distinction along the radial axis: pyramidal cells located closer to the stratum oriens (or ‘‘deep’’ sublayer) and stratum radiatum (or ‘‘superficial’’ sublayer) are different in size and density across a wide range of species.^47,49^ Numerous recent studies have validated this approximation, based on gene expression, anatomical features, afferent/efferent connections, biophysical properties, differential pyramidal-interneuron interactions, population cooperativity and behavioral correlates.^15,17–20,37,50–55^

Our birthdating observations based on in-utero electroporation support and refine the deep-to-superficial organization of subcircuits within the CA1 pyramidal layer. A subset of physiological parameters we examined varied monotonically with birthdate and were largely consistent with reported differences between deep and superficial cells. Average firing rates and afferent response properties of pyramidal neurons, as reflected by their theta phase shift during REM sleep, faithfully reflected the inside-out addition of newborn cells to the pyramidal layer (**Fig. 1**). However, in most measures – burstiness, SPW-R related firing rates, theta depth modulation, and spatial information content (**Fig. 1,3,6**) - early and late born neurons (deep and superficial, respectively) displayed quantitative similarities. These observations resonate with a recent report demonstrating that many biophysical and anatomical properties of ventral CA1 pyramidal neurons are better explained by birthdate than anatomical positioning.^33^ A small group of earliest born (E12.5) “pioneer” neurons exhibited unique biophysical properties and a broad dispersion in the pyramidal layer, with a tendency bias towards the stratum oriens.^33^ Taken together, these findings suggest that the deep-superficial distribution is graded rather than step-like, with neurons born at different time points showing broad and strongly overlapping topography (**Fig. 1**). Because of the anatomical overlap of different day-born neurons, the depth criterion alone is not a robust separator of either anatomical or physiological features.

### Neurons that are born together, wire and fire together

A dominant framework portrays neural systems in which sensory inputs sculpt connectivity and, consequently, dynamical patterns, assisted by plasticity (“neurons that fire together wire together”).^56^ In such models, there is a fundamental tension between competing features of plasticity and circuit stability, often leading to ‘catastrophic interference’ with previously stored knowledge.^57^ Alternatively, neuronal circuits can give rise to and maintain stable preconfigured dynamics, often referred to as attractors, manifolds, or neural schemata.^58–60^ Such preconfigured networks internally generate and maintain a large reservoir of activity patterns available for matching with novel experiences.^23,27,28,31,61–63^ Our study was designed to examine the potential source of such preconfigured dynamics.

The most robust finding of our experiment is a shared relationship among neurons born on the same day compared to neurons born on different days. Hippocampal neurons, born and developed together innervated the same target interneurons (**Fig. 2**). In complementary fashion, other studies have shown that same birthdate neurons exhibit structured connectivity between subregions of the hippocampus,^36^ with highly clustered connectivity onto specific dendritic compartments.^64^ Furthermore, CA1 pyramidal “sister” neurons born from the same parent progenitor receive common synaptic inputs from nearby fast spiking interneurons.^45^

Altogether, this wiring logic is reminiscent of microcircuits in the neocortex, where layer-specific neurons receive structured input and exhibit reciprocal connections between principal cells and interneurons.^65^ Furthermore, some interneuron types in superficial layers asymmetrically suppress spiking activity in deeper layers, routing firing patterns in a top-down direction.^66,67^ In the hippocampus, deep CA1 pyramidal cells receive stronger inhibition from parvalbumin-expressing basket cells, while superficial pyramidal cells provide stronger excitatory inputs to basket cells. Furthermore, within the deep sublayer, parvalbumin-expressing basket cells asymmetrically inhibit pyramidal cells with different target projections,^17,20^ allowing selective and differential routing of CA1 outputs. Thus, the fundamental motif organization rules may be similar in the hippocampus and isocortex.

The CA1 microcircuit domains, determined by embryonic development, constrained and predicted single neuron and population firing features in the adult brain. Same day-born neurons showed stronger spike correlations, firing together in the same sharp wave-ripples and the same theta cycles (**Fig. 3, 6, 7**). We also demonstrated that same day-born pyramidal neurons displayed overlapping spatial receptive fields and their population activity persisted in different environments, as probed by their preferential co-firing in the left and right arms of the maze (**Fig. 6**). Given their overlapping efferents outside the hippocampus,^33^ groups of same birthdate pyramidal neurons are likely to possess a unique capacity to discharge their postsynaptic targets across a wide range of brain states. Similar results may also hold for clonally related pairs of sister cells, which exhibit temporal cofiring in the hippocampal slice preparation,^45^ and are known to exhibit similar orientation tuning in the visual cortex.^68,69^

These findings support the preconfigured model and are corroborated by previous observations. In intracellular studies, spatially uniform depolarization of pyramidal cells gave rise to reliable firing at a fixed spatial position.^25,26^ Subsequent experiments used short-pulse optogenetic probing and artificial place field induction to reveal that induced fields preexisted where place fields with suprathreshold spiking subsequently emerged, and the locations of unmasked place fields were predictable from their correlated activity with peer neurons during sleep prior to optogenetic unmasking.^29,70^ In another study, firing rates of neurons during optogenetically induced and naturally occurring SPW-Rs were correlated, further illustrating circuit property constraints.^71^ Such preexisting circuit constraints can explain why place cell sequences in a novel environment can be predicted from the sharp wave ripple-associated “preplay” sequences during sleep prior to the experience.^27,28,30,72^ In developing rats, the first exploratory trip beyond the confines of the nest takes place between postnatal day 15 to 17, and CA1 pyramidal neurons already display adult-like place fields and ordered place cell sequences on the very first exploration.^73–75^ Our findings indicate that prenatal development of hippocampal circuits plays a critical role in forming a reservoir of preconfigured activity patterns that can be linked to novel experiences. In support of this pattern-matching hypothesis, theoretical work suggests that hippocampal pyramidal neurons can support a vast repertoire of place field arrangements in the absence of sensory cues given structured synaptic inputs from grid cells.^76^ Furthermore, learning to generate sequences over latent states has recently been shown to capture an impressive range of cognitive map and remapping phenomena.^77^

Same birthdate pyramidal neurons with shared spatial properties were often nearby neighbors. The existence of these functional microdomains is at variance with a strictly random organization view of spatial representation by hippocampal neurons,^4^ which has remained the prevailing view despite observations of clustered place cells using calcium imaging and early gene expression techniques.^78,79^ A practical explanation of this controversy is that tetrode recordings cannot reliably resolve place cells with overlapping place fields on the same tetrode and neighboring tetrodes are typically placed >400 µm from each other, precluding the detection of such domains. Furthermore, the difficulty of establishing a consensus on such peer domains might be explained by unique prenatal migration mechanisms in the hippocampus. In the neocortex, radially migrating excitatory neurons derived from the same progenitors migrate radially and organize themselves into vertical functional columns.^80^ In contrast, post-mitotic hippocampal neurons use multiple radial glial fibers as scaffolds and migrate radial-tangentially in a zigzag manner to establish horizontal clusters.^45,81^ Consistent with this generative process, same birthdate pyramidal neurons expressed overlapping spatial representations at anatomical distances ≥ 250 µm (**Fig. 6**), thus revealing a developmentally established functional microtopography in the hippocampus. Intermingling of different birthdate neurons in the ∼50 µm thin pyramidal layer may further clarify why the anatomical clustering of functional features has been hard to establish conclusively in the hippocampus. In future works, transcriptional profiling identifying novel marker genes of developmentally defined hippocampal neurons and their targeted perturbations (e.g., CRISPR–Cas gene editing) will be invaluable for causally testing the behavioral functions of microdomain organization.

### Wave of neurogenesis affects local connectivity and functional properties of pyramidal cells

A number of physiological measures (i.e., bursting, sharp wave-ripple related firing, theta depth modulation, and spatial information content) displayed a U-or inverted U-shaped relationship with birthdate. This profile reflects the known bell-shaped wave of neurogenesis,^35,36^ which was apparent in our data in the fractions of light responsive pyramidal neurons observed at four distinct birthdates (**Fig. 1**). Perhaps most notably, the expression rate of assemblies also took on a bell-shaped profile as a function of the birthdate of assembly members (**Fig. 4**). While intrinsic properties of different birthdate hippocampal neurons have recently been documented,^33^ the consequences of the wave-like neurogenesis profile on network-wide activity have not been considered by previous work.

Development of large networks is governed by robust self-organizing phenomena that go beyond the particulars of the individual constituents. The two key features of such networks with high degree of self-organization are growth and preferential attachment.^82^ Similarly, we found that the pace of neurogenesis alone cannot explain our observations, in that the bell-shaped assembly expression rates (**Fig. 4**) are not a simple consequence of the bell-shaped neurogenesis wave. Instead, our model (**Fig. 5**) suggested that, in addition to the neurogenesis wave (growth), a spike correlation rule (attachment) must also be in place to accommodate the empirically observed assembly behavior. This additional rule might be the birth rate-dependent correlated strengthening of the reciprocal connections between same day-born pyramidal cells and their interneurons^45^ or preferred co-innervation of same day-born neurons from upstream same day-born peer neurons.^36,64^ We further hypothesize that the U-shaped profile observed in a number of physiological parameters might reflect a fingerprint of this network-wide effect.

In summary, we suggest that the radial topography and heterogeneity of functional features within the CA1 pyramidal layer results from a combination of neuronal birthdate and the rate of neurogenesis, and these rules may generalize to other cortical networks.

## AUTHOR CONTRIBUTIONS

RH, HB, and GB designed the study. RH and YZ performed the experiments, and RH performed data analysis and modeling with help from YZ in analyzing histological data. RH and GB wrote the manuscript with contributions from other authors.

## ACKNOWLEDGEMENTS

The authors would like to thank Antonio Fernandez-Ruiz, Dhananjay Huilgol, Kathryn McClain, Sam McKenzie, Andrea Navas-Olive, Noam Nitzan, Manuel Valero, and Ipshita Zutshi for feedback on an early version of the manuscript. We also thank Xiaohan Wang, Takashi Yamaguchi and Jeremy Dasen for technical support. This work was supported by U19NS104590-01.

## LEAD CONTACT

Further information and requests for resources and data should be directed to the Lead Contact, György Buzsáki.

## DATA AND SOFTWARE AVAILABILITY

Custom lab software is available at https://github.com/buzsakilab/buzcode. Custom scripts for the current project will be made available upon request. The dataset generated for the current study will be made publicly available in the Buzsáki lab repository (https://buzsakilab.nyumc.org/datasets/).

## METHODS

### Animals

All experiments were conducted in accordance with the Institutional Animal Care and Use Committee of New York University Medical Center. Time-pregnant C57BL/6J female mice were either bred in-house or obtained from Charles River Laboratory. Timed pregnancies were prepared by co-housing males and females shortly before the dark cycle. Early morning of the next day was considered embryonic (E) age 0.5. Time pregnant mice and their offspring were kept on a regular light-dark cycle. Electrode-implanted adult mice (3-6 months) were housed individually on a reverse light-dark cycle.

### Histology

Animals were deeply anesthetized with isoflurane, and transcardially perfused with 4% paraformaldehyde. Brains were post-fixed in 4% paraformaldehyde for 1 day, and then moved to 1X PBS. Brain slices were taken at 50-75 μm (LEICA VT10000 S). Slices were mounted onto glass slides, stained with DAPI (SouthernBiotech), and imaged under a fluorescent microscope (OLYMPUS BX61VS) with a magnification of 10X under three filters: DAPI, TRITC (for tdTomato), and FITC (for EYFP).

### Quantification of normalized cell depth

CA1 pyramidal neuron coordinates were identified by thresholding expression in the red spectrum (tdTomato), followed by manual curation (ImageJ). In each section, the CA1 region was delimited according to the Mouse Brain Atlas.^83^The border between stratum pyramidale (SP) and stratum radiatum (SR) as well as the border between SP and stratum oriens (SO) were identified manually assisted by the high contrast of DAPI expression (**Supplementary Fig. 1**). Using custom written scripts, a perpendicular projection was made from each identified pyramidal neuron to the SP-SR border. Connecting the cell and its projection foot on the SP-SR border, an extension was made to cross the SP-SO border. This distance was taken to be the local thickness of the pyramidal layer. The perpendicular distance from each neuron to its projection onto the SP-SR border was divided by the local layer thickness to obtain the neuron’s normalized depth.^84^ n=3 E13.5 brains, n=4 E14.5 brains, n=3 E15.5 brains, and n=4 E16.5 brains were used for this quantification.

### In-utero hippocampal electroporation

Birthdating of CA1 pyramidal neurons was performed via in-utero electroporation^34^ in embryos of time pregnant females at E13.5 (n=26 mothers, ∼7% successful), E14.5 (n=4, ∼50% successful), E15.5 (n=13, ∼70% successful), and E16.5 (n=3, 100% successful). Success rate refers to the fraction of litters in which at least one offspring survived and expressed the electroporated DNA constructs. All surgeries were performed in the morning. Time-pregnant female mice were anesthetized with isoflurane (1.5-2% isoflurane at 1 L/min air flow rate) with vital signs monitored throughout the procedure. A subcutaneous injection of 5 mg/kg meloxicam was delivered for analgesia. The central portion of the uterine horn was extracted and placed on sterile gauze humidified with PBS. Using a pulled glass capillary, a plasmid DNA solution mixed with fast green FCF dye (Fisher) was injected (Picospritzer II, Parker) into the lateral ventricles of each embryo. The DNA solution consisted of ChR2-EYFP at 1.2 µg/µl (pAAV-CaMKIIa-hChR2(H134R)-EYFP; Addgene plasmid # 26969 ; http://n2t.net/addgene:26969 ; RRID:Addgene_26969) and pCAG-tdTomato at 0.65 µg/µl (Addgene plasmid # 83029 ; http://n2t.net/addgene:83029 ; RRID:Addgene_83029). Plasmids and fast green were diluted in sterile 1X PBS. Following injection, plasmids were electroporated towards the progenitors of CA1 pyramidal neurons using the triple electrode technique (CUY700 and CUY650 electrode series from Nepagene) (**Fig. 1A**).^34^ Electroporation involved 5 50ms pulses at 35-45 V (depending on the age of embryos), with 500ms interpulse intervals (ECM 830, Harvard Apparatus). Because the locus of targeted pyramidal neurogenesis (ventricular zone) occurs far away from the locus of interneuronal neurogenesis (ganglionic eminences), the probability of off-target expression in interneurons is very low. After the procedure, the uterine horn was placed back in the abdomen, and the anterior muscle wall and overlying skin were sutured. After birth, dams were cohoused with the pups until these have reached weaning age (3-4 weeks). At that point, the male and female pups were housed separately.

Using in-utero electroporation to measure the rate of neurogenesis (as is stated in the main text) might be confounded by many factors, such as the anatomy of the embryo, efficiency of pulsing and the surface area of the electrode. Yet, previous studies that used BrdU labeling of dividing cells in the hippocampal CA1 and CA3 regions reported a bell-shaped wave of neurogenesis,^36^ which was consistent with previous studies using autoradiography.^35^ We observed a similarly shaped distribution using in-utero electroporation (**Fig. 1E**). Other experiments have shown that electroporated DNA is taken up by dividing cells most effectively when these are in the phase of the cell cycle (S-G2) associated with the most effective incorporation of BrdU.^85^ Altogether, these results support our use of the term ‘birthdating’ with in-utero electroporation.

### Silicon probes coupled with optic fibers for optogenetics

High-density silicon probes were mounted on fully recoverable and reusable 3D printed micro drives.^86^ Silicon probes used in this study were the following: ASSY-156-E-1 (Cambridge NeuroTech), ASSY Int128-P64-1D (Diagnostic Biochips), and ASSY Int64-P32-1D (Diagnostic Biochips). The latter two types were dual-sided designs, equipped with recording sites on both sides of each shank. The spacing between shanks was 250 µm. Once mounted on a microdrive, probes were coupled with a single 100 µm diameter optic fiber with a ferrule attached to one of its ends (Thor labs). Blue light (450nm) was delivered via laser diodes (PL450B, Arrow Inc.) and controlled by the open-source Cyclops LED driver (https://github.com/jonnew/cyclops).^87^ Prior to coupling with the probe and implantation, optic fiber quality was assessed by measuring the maximum light intensity (PM100D, Thor labs) at the fiber tip. A maximum output of at least 8 mW/mm^2^ was required.

### Implantation and recording

Chronic recordings were performed from n=14 freely moving adult mice (10 males, 4 females; 3-6 months old; electroporated at E13.5, n=3; E14.5, n=3; E15.5, n=5; E16.5, n=3) for ∼4-12 weeks. The mice were anesthetized with 1.5–2% isoflurane and implanted with silicon probes coupled with optic fibers directed at the dorsal CA1 region. A stainless-steel screw was placed over the cerebellum for grounding and reference, and a craniotomy was drilled at -2mm A.P. and 1.7mm M.L. An acute probe equipped with an optic fiber was slowly descended above the hippocampus (∼600-1000µm), and square light pulses (100ms) were delivered to verify ChR2 expression. If reliable optical responses were not observed in either hemisphere, surgery was terminated, and the animal was euthanized with an overdose of isoflurane. Otherwise, a silicon probe / optic fiber attached to a 3D printed microdrive was implanted at a 45° angle along the anterior-posterior axis at a depth of approximately 700 µm. Due to lateralization of electroporated ChR2, approximately half of the animals were implanted in their left hemisphere and the remaining half in their right. Craniotomies were sealed with a mix of dental wax and mineral oil, and a copper mesh cage was constructed to provide electrical and mechanical shielding. Post operatively, animals received a single intramuscular injection of 0.06 mg/kg buprenorphine (0.015 mg/ml), and as needed for the following 1–3 days. Animals were allowed a 7-day recovery period before the start of experiments.

Following a 7-day recovery period, neural signals were recorded in the homecage while probes were descended into the CA1 pyramidal layer, which was identified physiologically via the sharp wave polarity reversal. Neural data was amplified and digitized at 30 kHz using Intan amplifier boards (RHD2132/RHD2000 Evaluation System, Intan). All recordings (92 sessions ranging in duration from 113.7 min to 475.6 min; median duration = 309.1 min) included a homecage period of sleep and wake.

### Behavior

A subset of n=8 mice (E13.5, n=1; E14.5, n=2; E15.5, n=2; E16.5, n=3) was trained on a spatial alternation task in a figure-8 maze. Animals were water restricted before the start of experiments and habituated to a custom-built 79×79cm figure-8 maze (**Fig. 6A**) raised 61cm above the ground.^88^ Over 3-5 days following the start of water deprivation, animals were shaped to alternate arms between trials to receive a water reward in the first corner reached after making a correct left/right turn. A 5-s delay in the start area was introduced between trials. The length of each trial was 205 cm. IR sensors were used to detect the animal’s progression through the task, and 3D printed doors mounted to servo motors were opened/closed to prevent the mice from backtracking (**Fig. 6A**). IR sensors and servo motors were controlled by a custom Arduino-based circuit.^88^ The position of head-mounted red LEDs was tracked with an overhead Basler camera (acA1300-60 gmNIR, Graftek Imaging) at a frame rate of 30Hz, and tracking data was aligned to the recording via TTL pulses from the camera, as well as a slow pulsing LED located outside of the maze. Animals were required to run at least 10 trials along each arm (at least 20 trials total) within each session. In all sessions that included behavior, animals spent ∼120min in the homecage prior to running on the maze, and another ∼120 minutes in the homecage after. All behavioral sessions were performed in the mornings.

### Stimulation protocol

Square light pulses in blocks of 5 with increasing light intensity and 200ms interpulse intervals were delivered to induce spiking in ChR2 expressing, birthdated pyramidal neurons. Pulse duration was fixed within each session but varied across (range 1.5-3ms; median=2ms within session). A range of 300-1600 blocks per session was delivered (median=800 blocks per session). Optogenetic stimulation was delivered at the end of each session after the recordings were completed, while the animal rested in its homecage. Prior to each recording session, light intensities were calibrated to a level with observable spiking, but no LFP deflection that would reflect a population effect (**Supplementary Fig. 1**). Sharp onsets and offsets were associated with a photoelectric artefact that took the form of a spikelet. To prevent such artefacts from propagating to spike sorting and unit identification, raw data was clipped out in the interval shortly before the onset (0.15ms) and after the offset (0.8ms) of each brief pulse.

## QUANTIFICATION AND STATISTICAL ANALYSIS

### Unit isolation and classification

Spikes were extracted and classified into putative single units using KiloSort1.^89^ Manual curation was performed in the Phy2 software with the aid of custom-built plugins (https://github.com/petersenpeter/phy2-plugins). Throughout the manual curation step, isolation quality was judged by inspecting cross-correlograms for incorrect splits of single units (i.e., auto-correlogram structure detectable in the cross-correlogram). Cells were classified as putative pyramidal cells and interneurons via CellExplorer (https://cellexplorer.org/pipeline/cell-type-classification).^90^ Briefly, putative interneurons were identified via hard thresholds imposed on the waveform shape (trough to peak), and the auto-correlogram rise and decay time constants.The dataset includes a total of 6,699 well-isolated putative pyramidal cells and 1,919 putative interneurons (1449 narrow waveform, 470 wide waveform; **Supplementary Fig. 2**).

### State scoring

State scoring was performed as described previously (https://github.com/buzsakilab/buzcode/blob/dev/detectors/detectStates/SleepScoreMaster/SleepScoreMaster.m).^91^ First, the local field potential (LFP) was extracted from wideband data by lowpass filtering (sinc filter with a 450 Hz cut-off band) and downsampling to 1250Hz. Three signals were used for state scoring: broadband LFP, narrowband theta frequency LFP and electromyogram (EMG). Spectrograms were computed from broadband LFP with FFT in 10s sliding windows (at 1s), and PCA was computed after a Z transform. The first PC reflected power in the low (< 20Hz) frequency range, with oppositely weighted power at higher (>32Hz) frequencies. Theta dominance was quantified as the ratio of powers in the 5–10Hz and 2–16Hz frequency bands. EMG was estimated as the zero-lag correlation between 300-600Hz filtered signals across recording sites. Soft sticky thresholds on these metrics were used to identify states. Briefly, high LFP PC1 and low EMG was taken to be NREM, high theta and low EMG was considered REM, and the remaining data was taken to reflect the waking state. All assignments were inspected visually and manually curated wherever appropriate (https://github.com/buzsakilab/buzcode/blob/dev/GUITools/TheStateEditor/TheStateEditor.m).

### SPW-R detection

SPW-Rs were detected as described previously^92^ from manually selected channels located in the center of the CA1 pyramidal layer (https://github.com/buzsakilab/buzcode/blob/master/detectors/detectEvents/bz_FindRipples.m). Broadband LFP was bandpass-filtered between 130 and 200 Hz using a fourth-order Chebyshev filter, and the normalized squared signal was calculated. SPW-R maxima were detected by thresholding the normalized squared signal at 5×SDs above the mean, and the surrounding SPW-R start/stop times were identified as crossings of 2×SDs around this peak. SPW-R duration limits were set to be between 20 and 200 ms. An exclusion criterion was provided by designating a ‘noise’ channel (no detectable SPW-Rs in the LFP), and events detected on this channel were interpreted as false positives (e.g., EMG artifacts).

### Optogenetic tagging of birthdated pyramidal neurons

ChR2 expressing neurons have been shown to fire at characteristic latencies with respect to stimulus onset.^93–96^ However, since this may also be true of some non-ChR2 expressing neurons due to polysynaptic effects (e.g., precisely timed rebound from inhibition), we additionally tested the reliability of firing following stimulus onset as opposed to preceding it. Because interneurons were expected to respond due to strong convergent inputs from their presynaptic pyramidal neurons,^40^ only pyramidal neurons identified via unit classification were considered for optogenetic tagging.

The latency to spike effect was quantified using SALT.^97^ For each pyramidal neuron, the distribution of latencies to first spike in 10ms windows following each pulse was compared to independent “baseline” latency to spike distributions (n=200) with respect to random timepoints outside of optogenetic stimulation. To avoid the effect of slowly changing firing rates on spike latencies, the random timepoints were selected in a period before the first pulse onset in an interval of equal duration to that between the first and last pulse. A p-value was obtained by comparing the median distance (Jensen-Shannon divergence) between the post-stimulus and baseline distributions against a null computed from distances between baseline distributions. P<=0.001 was considered significant.

To test the reliability of poststimulus firing, we adapted a routine for the detection of monosynaptic connections.^40,98^ As described previously, we computed the peristimulus time histogram (1ms bins) and smoothed with a hollowed Gaussian (15ms SD) to obtain a baseline estimate of persistimulus firing matched for slow changes in firing rate. We considered spike counts in three time bins surrounding that of peak poststimulus firing and assessed whether any of these were significantly greater than baseline counts, assuming a Poisson distribution over counts at each bin. Furthermore, the bin of maximum poststimulus firing was compared with bins at similar lags preceding pulse onset, to test whether peak firing following the stimulus was significantly greater than firing preceding the stimulus. In each case, alpha was set at 0.001, and Bonferroni corrected for multiple comparisons.

All the above-described tests needed to be passed for a pyramidal neuron to be considered optogenetically tagged.

### Monosynaptic connection analyses

Cross correlograms (CCGs) between pairs of neurons were constructed (0.8ms bins) from spikes occurring outside of optogenetic stimulation. Statistical detection of monosynaptic connections was broadly similar to the reliability test described above for optogenetic tagging. For full details of the algorithm, see English et al., 2017.^40^ Only connections from pyramidal neurons to interneurons were considered further.

To assess the effective strength of synaptic coupling at different presynaptic firing rates, the spike transmission probability metric was computed. First, CCGs were constructed from subsets of pyramidal spikes whose preceding interspike intervals fell in a specific range.^40^ The resulting CCG spike counts were divided by the number of presynaptic spikes in the subset to obtain the probability of interneuron firing at various lags with respect to the presynaptic spike. Values at 0.8-2.8ms lags that exceeded the baseline probability of postsynaptic firing (obtained by convolving CCGs with an 8ms SD hollowed Gaussian) were integrated, resulting in spike transmission probability.

Convergence onto interneurons was assessed for pairs of pyramidal neurons. For each pair, the number of interneurons they both projected to (i.e., convergence) was divided by the number of interneurons they targeted collectively, resulting in a convergence index that took on values between 0 and 1.

### Cofiring analysis

The spike count in each interval (either a SPW-R, or a theta cycle) was computed, resulting in a vector of spike counts for each pyramidal neuron. Pearson correlation coefficients between spike count vectors of different neurons were computed to estimate the SPW-R and theta cycle related co-firing of each pair.

### Same versus different birthdate pairs

All unique pairs of optogenetically tagged pyramidal neurons in a given recording session were considered to be of the same birthdate (SBD). This group was compared to one comprising of all unique pairs in which one pyramidal neuron was optogenetically tagged and the other was not. These pairs were considered of different birthdate (DBD). It is possible that some untagged neurons failed to be targeted by in-utero electroporation, and therefore a fraction of our “DBD group” could contain some SBD pairs. However, we assumed this fraction would be sufficiently small given the amount of true DBD neurons and continued to use the term “DBD” for convenience.

### Theta cycle detection

Because theta phase shifts along the radial axis,^99^ a channel with a positive sharp wave (i.e., above the center of the pyramidal layer) was selected to ensure consistency of extracted phases across recordings. Broadband LFP was bandpass-filtered between 6 and 12 Hz using a fourth order Chebyshev filter. The Hilbert transform of the filtered signal was computed, and its absolute value and angle at each timepoint were taken to be the theta band amplitude and phase, respectively. Intervals with theta band amplitude 1 SD above the mean were considered for theta cycle detection. Within these intervals, timepoints where the phase crossed 0° were identified as peaks of theta, and timepoints of consecutive theta peaks were considered the onsets and offsets of individual theta cycles (all throughout, peaks are at 0° and 360° and troughs at 180°). Only theta cycles occurring within identified waking periods (see State scoring) were considered for analysis.

### REM shifting and theta depth modulation

Theta phase was extracted in the waking and REM periods identified via state scoring. Broadband LFP was filtered, and Hilbert transformed to extract amplitude and phase, as described above. In the waking state, intervals with theta band amplitude above the mean were considered, while REM periods were considered in their entirety, since high theta amplitude was required for their detection in the first place. Each spike time falling within these periods was assigned a theta phase, resulting in a separate phase distribution in wake and REM. Phase locking was tested in each distribution (Rayleigh test), and when significant (p<0.01), the mean direction and mean resultant length of the circular phase distribution were taken as the preferred theta phase and theta depth modulation, respectively. Pyramidal neurons that were significantly phase locked in wake and REM were designated as REM shifting if their preferred phase was between 120° and 300° during wake, and outside this interval during REM.^18^

### ICA assembly analysis

To detect assemblies reflecting higher order coactivity among pyramidal neurons, we performed independent component analysis (ICA) as has been described previously.^43,44,100^ Spikes from pyramidal neurons recorded in the homecage were binned at 1ms resolution, smoothed with a Gaussian (25ms FWHM), and the resulting timeseries was z scored. The number of assemblies was based on the N principal components whose variances exceeded an analytical threshold based on the Marcenko-Pastur distribution describing variances expected for uncorrelated data. The z-scored matrix was projected into the subspace spanned by these N components, and ICA was performed to extract assemblies (each corresponds to an IC). As both the scale and sign of IC weights are arbitrary, IC weights were rescaled to unit norm and multiplied by the sign of the highest absolute value weight. Pyramidal neurons whose IC weights exceeded 2 standard deviations above the mean weight were considered “assembly members”, and all other neurons as “assembly non-members”. Only assemblies with at least one optically tagged (i.e., birthdated) assembly member were considered for further analysis. The expression strength of each assembly was computed as

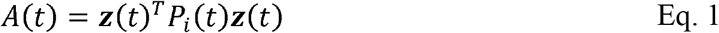

where *P*_*i*_(*t*)is the projection matrix (outer product, diagonal set to zero) of the i-th IC. *A*(*t*)quantifies the moment-to-moment assembly expression strength as the similarity between an IC and the instantaneous, z-scored firing pattern recorded across recorded pyramidal neurons (i.e., a projected z-score). Peaks that exceeded 2 standard deviations above the mean expression strength were taken as timepoints of assembly expression, which were used for subsequent analysis (e.g., assembly expression rates in SPW-Rs).

For the assembly analyses in **Fig. 7**, spikes were binned at 25ms resolution and z scored. To detect assemblies associated with each held-out, birthdated pyramidal neuron, we identified bins that contained its spikes occurring in the period on the maze and performed ICA on the submatrix of remaining pyramidal neurons. To quantify the time-resolved expression of extracted assemblies, ICs were projected on a matrix that was binned finely (1 ms) and smoothed (25ms FWHM Gaussian) before z-scoring. In this case, all bins of the z-scored spike matrix were used. Timepoints of assembly expression and identities of assembly members were extracted as described above.

### Linear-nonlinear Poisson model for exploring assembly dynamics

At least two accounts could explain the qualitative similarity between assembly expression rates (**Fig. 4E**), and the bell-shaped fraction of neurons labeled at each birthdate (**Fig. 1E**). One explanation may be based on the observation that firing rates and participation in SPW-Rs also exhibited a bell-shaped relationship with birthdate. Differences in (SPW-R)-related firing across birthdates may bias correlations, which may affect (SPW-R)-related expression rates of the detected assemblies. An alternative explanation is that SBD neurons exhibit pairwise correlations over and above those expected from the observed firing rate differences. These excess correlations may then interact with the bell-shaped rate of neurogenesis to produce differences in assembly expression rates. To explore these two accounts, we generated spike trains with controlled firing rates and pairwise correlations (**Supplementary Fig. 7**),^46^ and compared their assembly expression rates to those observed in data.

We relied on a nonlinear transformation of multivariate Gaussian random variables to generate non-negative stochastic processes with controlled mean and covariance:

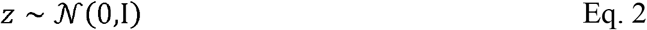

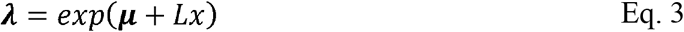

where *L* is a lower triangular matrix, such that Σ = *LL*^T^ and μ are the predefined covariance and mean of a Gaussian. An exponential transformation of samples from such a Gaussian results in a non-negative, log-normally distributed multivariate processλ, with the following mean and covariance:

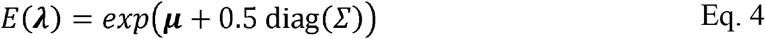

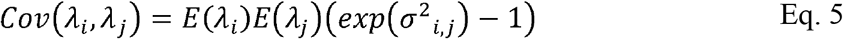

where diag (Σ)is a vector holding the Gaussian variances, and σ^2^_*ij*,_the Gaussian covariances. Each λ_*i*_ can be interpreted as the time varying firing rate of a neuron. By controlling the Gaussian variances diag (Σ)and means **μ**, we controlled both the firing rates λ_*i*_and their covariances Cov (λ_*i*_,λ_*i*_) Furthermore, (λ_*i*_,λ_*i*_) covariance depended on σ^2^_*i,j*_, which can be interpreted as an additional source of covariation that interacts with the joint firing rates of the individual neurons. Covariance Σ was generated using a squared exponential kernel:^101^

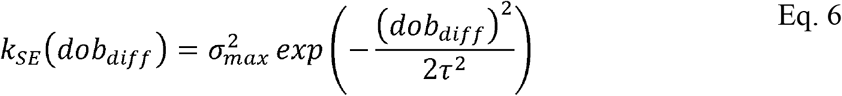

where *dob*_*diff*_refers to the difference of birthdates between two neurons. Maximum Gaussian covariance 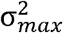 is achieved when |dob_*diff*_| is zero, and decays at a timescale τas dob_*diff*_ increases. This kernel function captures the notion that neurons born close together in time exhibited higher covariation in their firing.

The bell-shaped wave of neurogenesis was modeled as a Gaussian with mean 14.5 (denoting embryonic age), and standard deviation of 1 day. Each simulation began by drawing n=350 units with birthdates set according to this distribution. Gaussian covariance σ was specified following Eq. 6, and Eq. 4 was solved for **μ**, which was set to ensure that average firing rates followed those empirically observed in SPW-Rs (**Fig. 3D**). Specifically, firing rates of units <E14 were sampled from the E13.5 firing rate distribution, units ≥E14 and ≤E15 were drawn from the E14.5 distribution, units ≥E15 and ≤E16 from the E15.5 distribution, and units >E16 from the E16.5 distribution. Samples were generated according to Eq. 2 and 3 to generate non-negative rate functions λ_*i*_, and spike counts were simulated from the corresponding inhomogeneous Poisson processes. Each time step was assumed to be 25ms, and 10k samples (250s) were generated per simulation.

The resulting spike matrix was z-scored, and assemblies were detected and tracked using ICA as described above. For a given set of parameters, simulations were run until n=200 assemblies were extracted. Since each simulated neuron had a prespecified birthdate, the birthdate associated with each assembly was the average across assembly members (IC weights >2SD above the mean). To compare the birthdate-dependent profile of simulated assembly expressions to data, simulated expression rates were z-scored and the average across empirically observed assembly rates was added to each value. The shape of simulated assembly expressions as a function of birthdate was captured by fitting a Gaussian process nonlinear regression model (http://www.gaussianprocess.org/gpml/code/matlab/doc/),^101^ and the negative log predictive probability under this model was computed for the observed average assembly expression rate at each birthdate. The cumulative negative log probability was considered as the error.

Simulations were performed for a range of Gaussian covariances 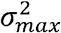and decay timescales τ to test the influence of firing rate differences (small 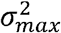) and the influence of additional correlations (large 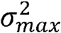) on the organization of assemblies. The simulation results in **Fig. 5D** were robust to the choice of above-described free parameters, as well as to the choice of error (e.g., MSE instead of log probability) for comparison with observed results (data not shown).

### Spatial ratemap analyses

Trials in the figure-8 maze were linearized, and velocity in each trial was estimated with a Kalman filter. Considering moments with speed >1.5cm/s, the number of spikes and time spent in each 1cm bin was computed separately for left and right trials and smoothed with a 9cm full width at half maximum Gaussian. The average ratemaps for left and right trials were computed as the smoothed spike counts normalized by the smoothed occupancy.

For spatial ratemap correlation between pyramidal neurons, ratemaps for left and right trials were concatenated, and the Pearson correlation coefficient was computed. The central stem portion (common between left and right trials) was excluded for the purpose of this analysis. Spatial information in bits/sec was computed as described previously.^102^

### Statistical analysis

Statistical tests involving multiple group comparison (i.e., ANOVA and Kruskal-Wallis) were performed non-parametrically with bootstrap resampling to generate null distributions of the relevant test statistic (i.e., F and H statistics, respectively). Specifically, datapoints across groups were pooled, and reassigned to groups randomly with replacement. N=5000 resamplings were performed to generate the null distributions. The same was performed for the q statistic for Tukey post-hoc comparison between groups. All post hoc comparisons were two tailed.

## SUPPLEMENTARY FIGURES

**Supplementary Figure 1.**
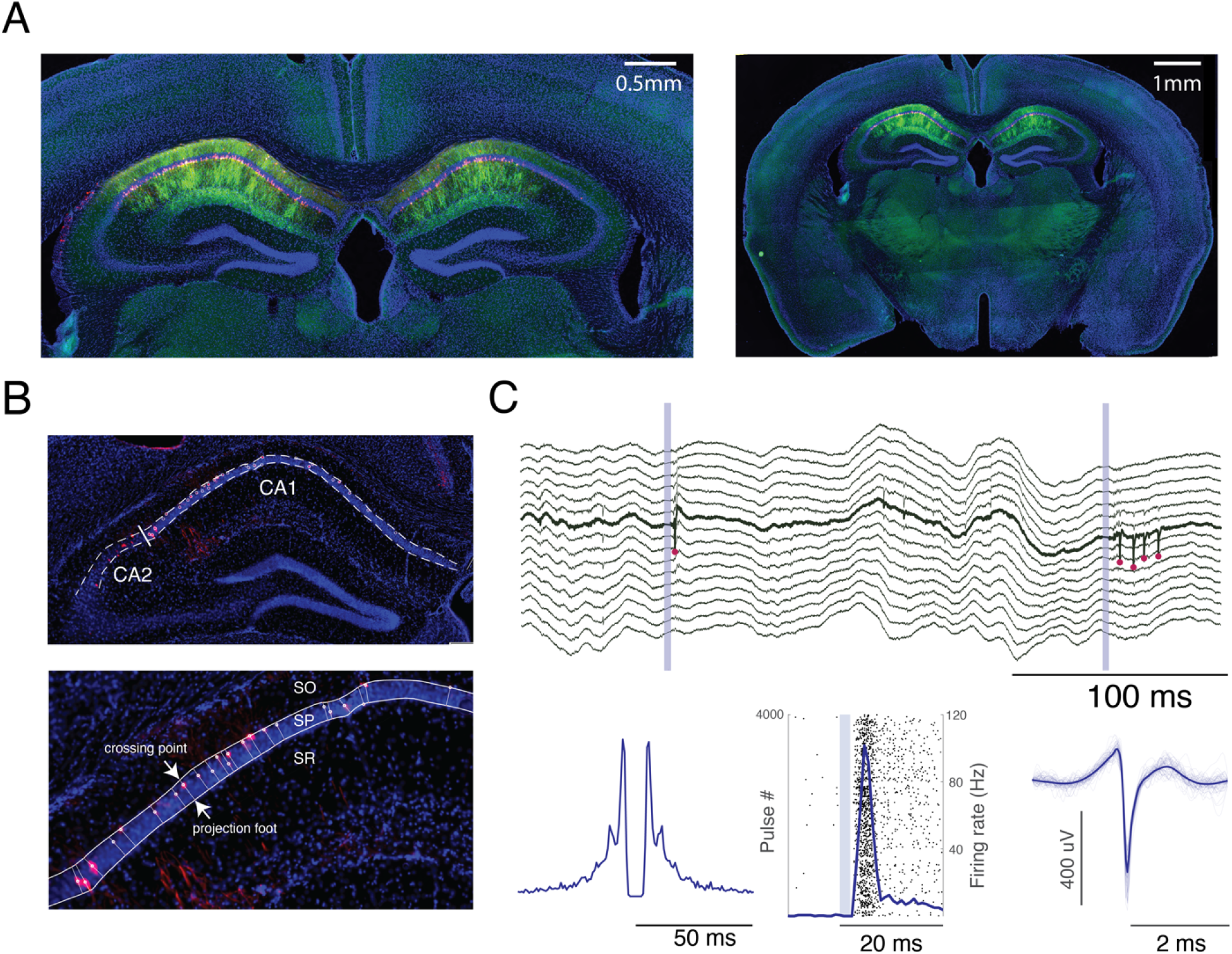
Sparse expression in CA1 following in-utero electroporation. (**A**) Left: Closeup of an example coronal section showing a typical expression profile of ChR2-EYFP (green) and tdTomato (red) resulting from in-utero electroporation at E15.5. Expression was restricted to the CA1 subregion of the hippocampus. Right: A full view of the section showing that expression was restricted to the hippocampus. (**B**) Top: Example histology section demonstrating confinement of expression to dorsal CA1 (different brain from that in (A)). Manually drawn borders (dashed) delimit the pyramidal layer. The thick white line illustrates the border between CA1 and CA2, which was approximated by comparing with the Mouse Brain Atlas. White circles are tdTomato+ puncta identified via ImageJ. Bottom: Closeup showing the projection foot of each tdTomato+ punctum on the border between str. pyramidale (SP) and str. radiatum (SR), from which each neuron’s radial depth was computed. Depth was normalized by the distance between the projection foot and crossing point with the border between SP and str. oriens (SO). (**C**) Top: Wideband (30 kHz) neural activity recorded on a single shank around the time of brief light pulses (2ms; blue). The bolded channel shows the spiking activity (red dots) of a pyramidal neuron that responded to blue light. Note single spike (first stimulus) and burst (second stimulus) responses. Bottom: Autocorrelogram (left) of the highlighted neuron’s spike train reveals a bursting firing profile, and the peristimulus time histogram (middle) demonstrates its fidelity of firing following light offset. Waveforms of spontaneous spikes are shown on the right (light blue, n=50 spikes; dark blue, average across all).

**Supplementary Figure 2.**
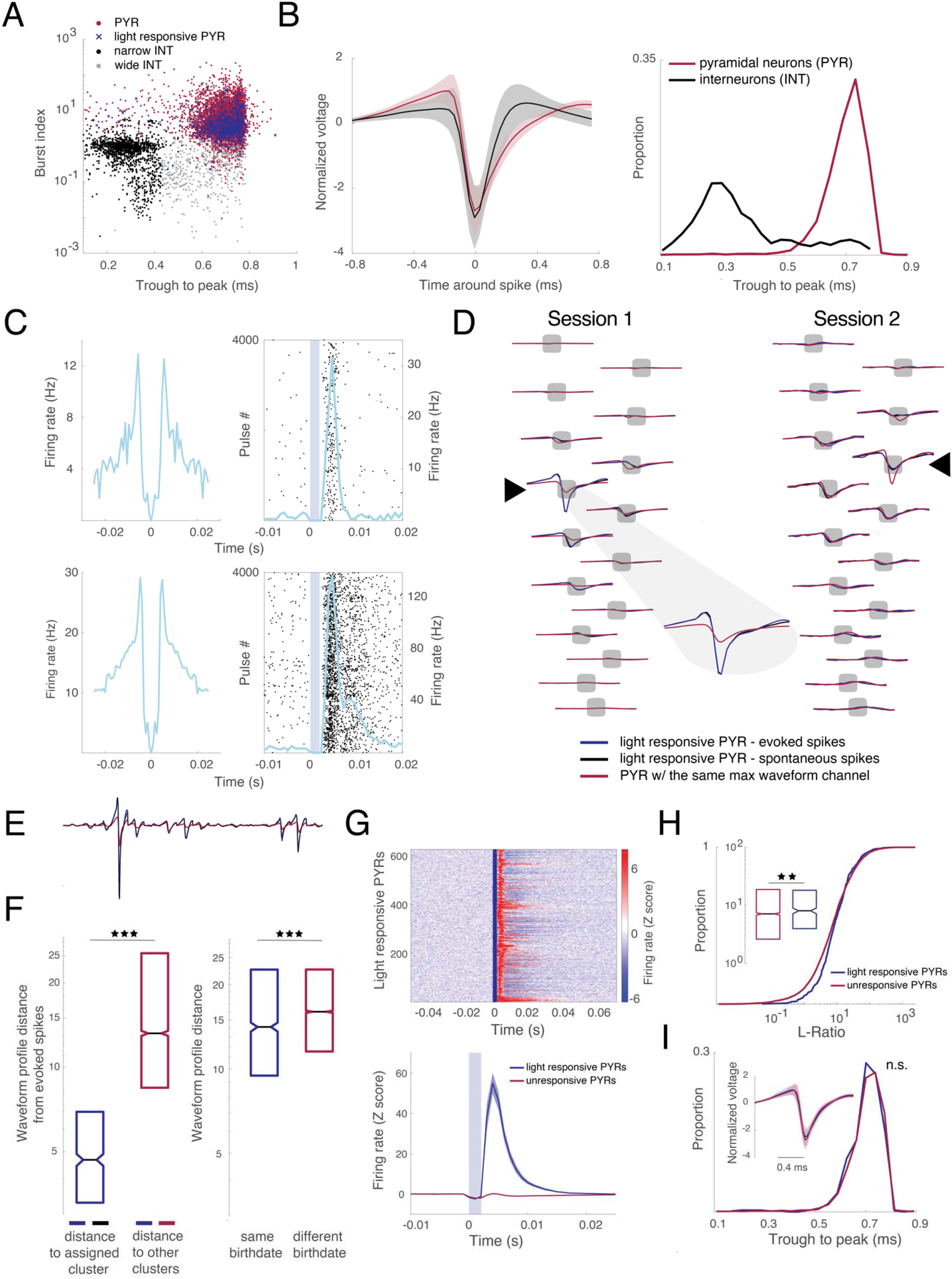
Neuron clustering, classification and optogenetic tagging of CA1 neurons. (**A**) Filtered waveform trough-to-peak (in ms) by burst index for each isolated unit (n=8619). Units are color coded to highlight putative pyramidal neurons (red, n=6699), narrow waveform interneurons (black, n=1449), and wide waveform interneurons (grey, n=470). Blue crosses highlight light-responsive pyramidal neurons (n=624, 9.31% of all pyramidal cells). (**B**) Left: Mean±SD of the filtered waveforms for all putative pyramidal neurons (red) and all interneurons (black). Right: Distribution of trough-to-peak values highlights the bimodality of waveform shapes used for cell classification. (**C**) Additional examples of light responsive pyramidal neurons, as in **Fig. 1D**. (**D**) Examples of unit separation on the same shank from two recording sessions. Average waveform profiles are shown across recording channels for two simultaneously recorded pyramidal neurons (light responsive, non-responsive) with maximum waveform amplitudes occurring on the same channel (highlighted with a black triangle). Blue, average of filtered waveforms of a light responsive pyramidal neuron from spikes occurring within 10ms of each brief light pulse. Black, average of filtered waveforms of the same light responsive pyramidal neuron from a random subset of 1000 spikes occurring outside of light pulses. Red, average of filtered waveforms of a non-responsive pyramidal neuron. In both examples, waveform profiles of evoked (blue) and spontaneous (black) spikes are indistinguishable (superimposed) and different from the waveform profile of the non-responsive (red) neuron whose maximum waveform occurred on the same channel. This is further highlighted in the blowup. (**E**) Waveforms from Session 1 in (D) (same color scheme), concatenated across channels to form a vector. The L2 norm of the difference between vectorized waveform profiles was computed to quantify the distance between waveform profiles. (**F**) Left: Waveform profile distances were computed between evoked and spontaneous spikes of light responsive pyramidal neurons (blue, median=4.78; n=624), and between evoked spikes of light responsive pyramidal neurons and spontaneous spikes of other pyramidal neurons whose maximum amplitude waveform occurred on the same channel (red, median=13.85; n=7879 neurons pairs; p=8.51e-248; Wilcoxon rank-sum test). Evoked spikes were most similar to the spikes of their assigned cluster, rather than to spikes of clusters of nearby neurons. This result argues against the possibility of spike sorting errors having significantly affected the optotagging of birthdated pyramidal neurons. Right: Waveform profile distances between all pairs of SBD (blue, median=14.12; n=1572) and DBD pyramidal neurons (red, median=16.007; n=14286; p=5.5748e-14; Wilcoxon rank-sum test) recorded on the same shank, irrespective of which channel. Only spontaneous spikes occurring outside of light stimulation were considered. Yet, pairs of SBD neurons had more similar waveform profiles than pairs of DBD neurons, possibly resulting from an effect of light-induced synchrony on template identification in Kilosort1. We performed resampling analyses to control for this possible confound of pairwise correlations (**Supplementary Fig. 6**). (**G**) Top: Z-scored PSTHs surrounding the onset of brief light pulses of pyramidal neurons that passed the statistical tests to be considered light responsive (n=624). Bottom: PSTH mean±S.E.M of light responsive pyramidal neurons (blue, n=624) and of non-responsive pyramidal neurons (n=6075). (**H**) Cluster isolation quality using the L-Ratio metric^103^ for light-responsive pyramidal neurons (blue, median=8.045; n=624) and non-responsive pyramidal neurons (red, median=7.1087; n=6075; p=0.0016; Wilcoxon rank-sum test). Clusters of light-responsive pyramidal cells were less well separated, leading us to perform resampling control analyses for the pairwise correlation results (**Supplementary Fig. 5**). (**I**) Waveform shape quantification using the trough to peak metric for light responsive (blue, mean=0.6946; n=624) and non-responsive pyramidal neurons (red, mean=0.6987; n=6075; p=0.1961; two-sample t-test). Inset: mean±SD of the filtered waveforms in each group.

**Supplementary Figure 3.**
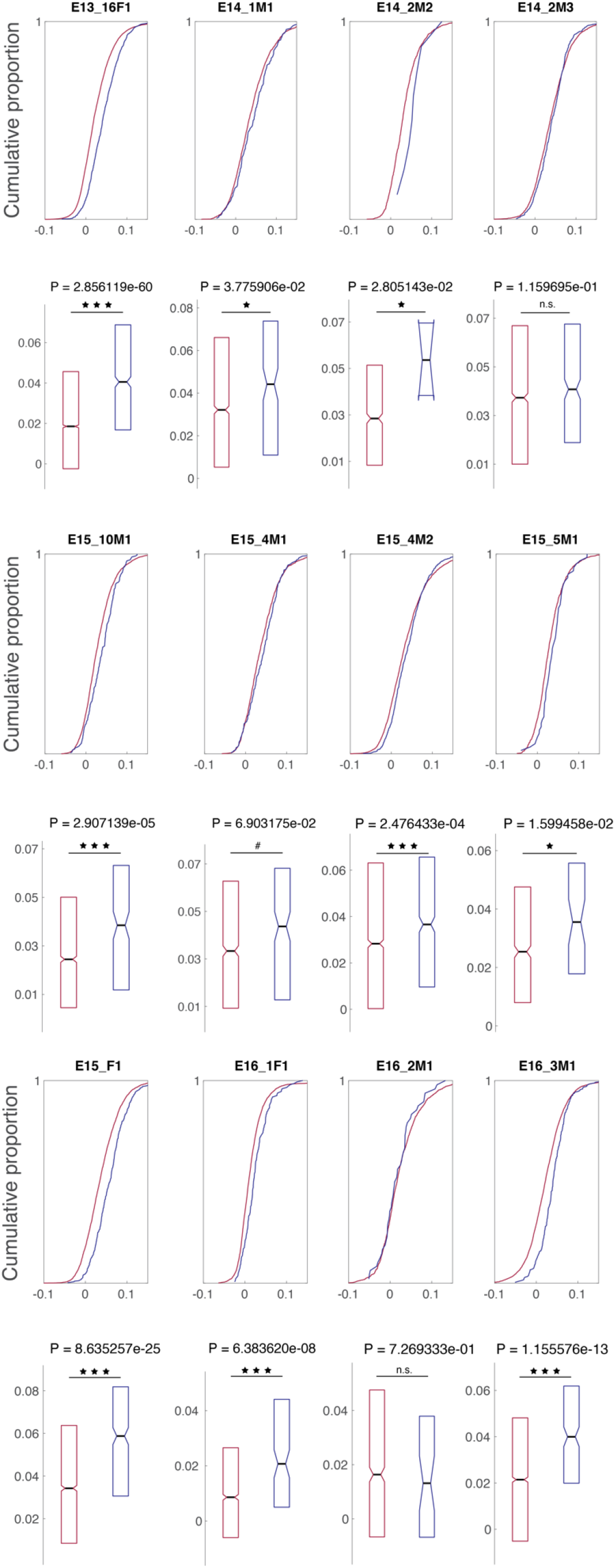
SPW-R correlations for pyramidal neuron pairs in individual mice. Cumulative distributions and box plot summaries of pairwise correlations in SPW-Rs for pairs of SBD (blue) and DBD (red) neurons in individual animals. SBD pyramidal neurons exhibited higher cofiring in SPW-Rs than DBD pyramidal neurons in 10/12 animals. P-values of Wilcoxon rank-sum tests are shown above each box plot summary. Two animals electroporated at E13.5 were excluded due to a lack of SBD pairs. In these two mice only a single light-responsive pyramidal neuron per session was recorded.

**Supplementary Figure 4.**
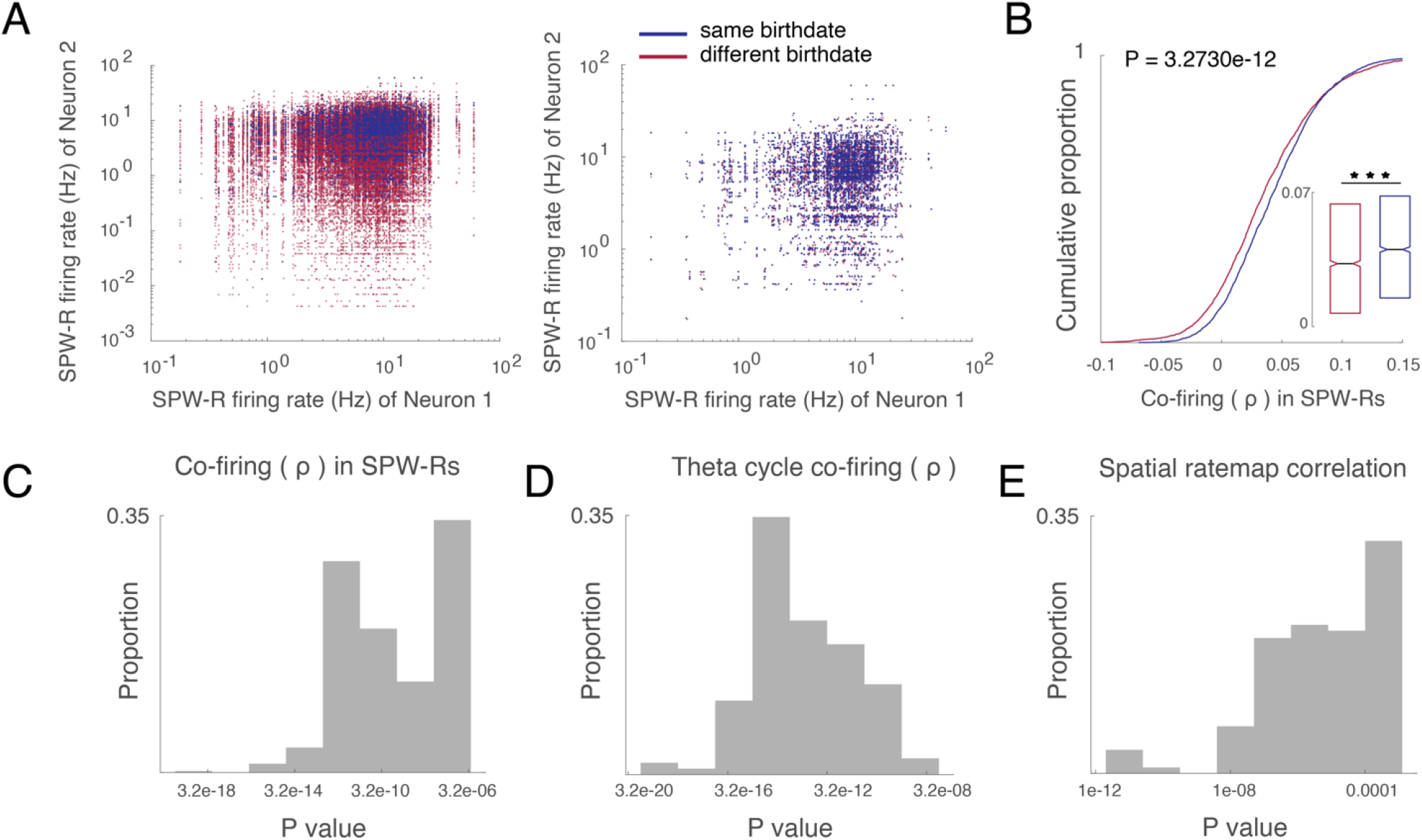
Resampling firing rate statistics support results in Fig. 3F and Fig. 6C,F. (**A**) Left: Joint firing rate distribution in SPW-Rs for all pairs of SBD (blue, n=3531) and DBD (red, n=44201) pyramidal neurons that went into the pairwise correlation analysis in **Fig. 3F**. Right: The joint firing rate distribution of DBD pairs was resampled according to the empirical joint probability mass function of joint firing rates of the SBD pairs. This procedure effectively matches the pairwise firing rate statistics and the group sizes. (**B**) Pairwise correlation in SPW-Rs for SBD (blue, n=3531) and resampled DBD pairs (red, n=3531; p=3.2730e-12, Wilcoxon rank-sum test). Note the persisting difference between the two groups. (**C**) Distribution of Wilcoxon rank-sum test p-values for the comparison in (B), following 500 independent resamplings of the DBD group. All p-values were well below the common significance level of 0.05. (**D**) Same as (A) to (C) for 500 resamplings of DBD joint firing rates during theta cycles, and the resulting comparisons between pairwise theta cycle correlations of SBD and resampled DBD pairs (as in **Fig. 6C**). (**E**) Same as (A) to (C) for 500 resamplings of DBD joint firing rates during theta cycles, and the resulting comparisons between spatial ratemap correlations of SBD and resampled DBD pairs (as in **Fig. 6F**). Altogether, these resampling results demonstrate that the differences shown in main **Fig. 3F** and **Fig. 6C**,**F** cannot be explained by difference of firing rates or by the different n’s in the compared groups.

**Supplementary Figure 5.**
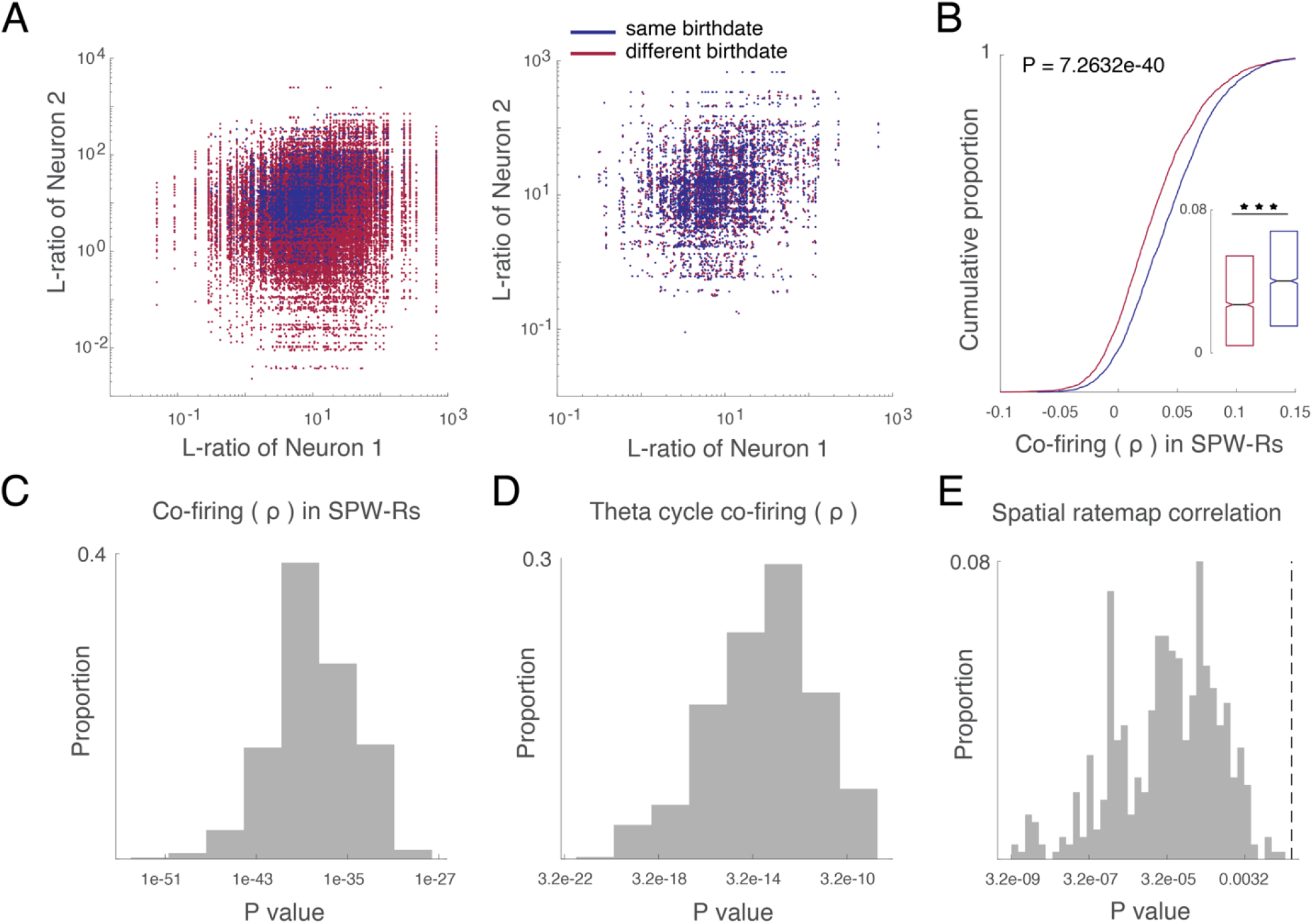
Resampling L-ratio statistics support results in Fig. 3F and Fig. 6C,F. (**A**) Left: Joint L-ratio distributions^103^ for all pairs of SBD (blue, n=3531) and DBD (red, n=44201) pyramidal neurons that went into the pairwise correlation analysis in **Fig. 3F**. Right: The joint L-ratio distribution of DBD pairs was resampled according to the empirical joint probability mass function of L-ratios of the SBD pairs. This procedure effectively matches the pairwise cluster isolation quality and the group sizes. (**B**) Pairwise correlations in SPW-Rs for SBD (blue, n=3531) and resampled DBD pairs (red, n=3531; p=7.2632e-40, Wilcoxon rank-sum test). (**C**) Distribution of Wilcoxon rank-sum test p-values for the comparison in (B), following 500 independent resamplings of the DBD group. All p-values were well below the common significance level of 0.05. (**D**) Same as (A) to (C) for 500 resamplings of DBD joint L-ratios, and the resulting comparisons between pairwise theta cycle correlations of SBD and resampled DBD pairs (as in **Fig. 6C**). (**E**) Same as (A) to (C) for 500 resamplings of DBD joint L-ratios during theta cycles, and the resulting comparisons between spatial ratemap overlap of SBD and resampled DBD pairs (as in **Fig. 6F**). Vertical dashed line indicates p=0.05.

**Supplementary Figure 6.**
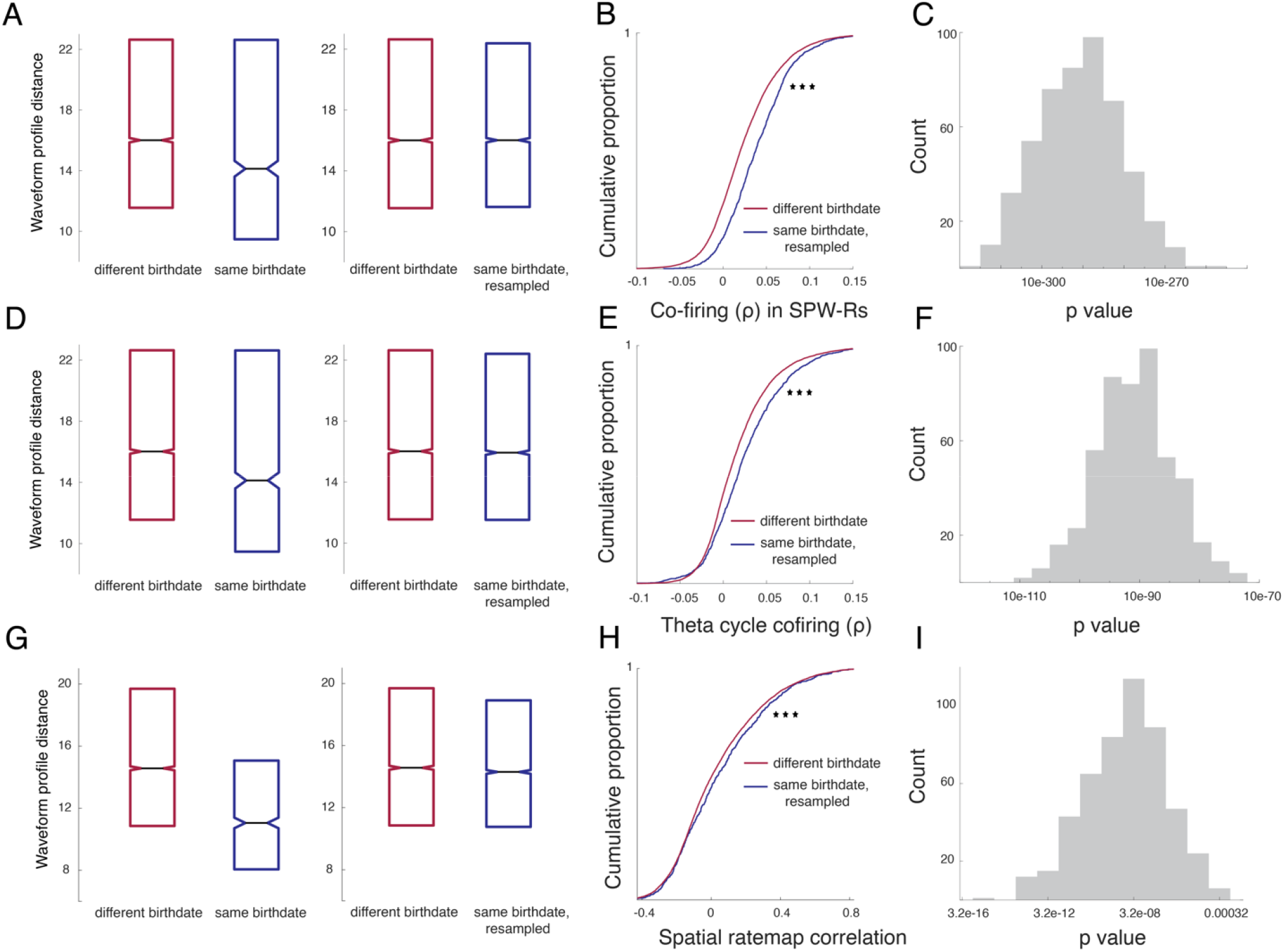
Resampling waveform profile statistics support results shown in main Fig. 3F and Fig. 6C,F. (**A**) Left: Waveform profile distances (as shown in **Supplementary Fig. 2F**) for all pairs of SBD (blue, n=1572) and DBD (red, n=14310) pyramidal neurons that were recorded on the same shank. Right: The waveform profile distance distribution of SBD pairs was resampled according to the empirical probability mass function of waveform profile distances of DBD pairs. This procedure effectively matches the statistics and groups sizes. Note that the SBD distribution was resampled since it was considered to be more affected by spike sorting errors, as reflected in lower waveform profile distances between same-shank SBD pairs. (**B**) Pairwise correlations in SPW-Rs for resampled SBD (blue, n=14164) and DBD pairs (red, n=14164; p=2.42e-274, Wilcoxon rank-sum test). (**C**) Distribution of Wilcoxon rank-sum test p-values for the comparison in (B), following 500 independent resamplings of the SBD group. All p-values were well below the common significance level of 0.05. (**D**)-(**F**) Same as (A) to (C) for 500 resamplings of SBD waveform profile distances, and the resulting comparisons between pairwise theta cycle correlations of DBD and resampled SBD pairs recorded on the same shank (as in **Fig. 6C**). (**G**)-(**I**) Same as (A) to (C) for 500 resamplings of SBD waveform profile distances, and the resulting comparisons between spatial ratemap overlap of DBD and resampled SBD pairs recorded on the same shank (as in **Fig. 6F**).

**Supplementary Figure 7.**
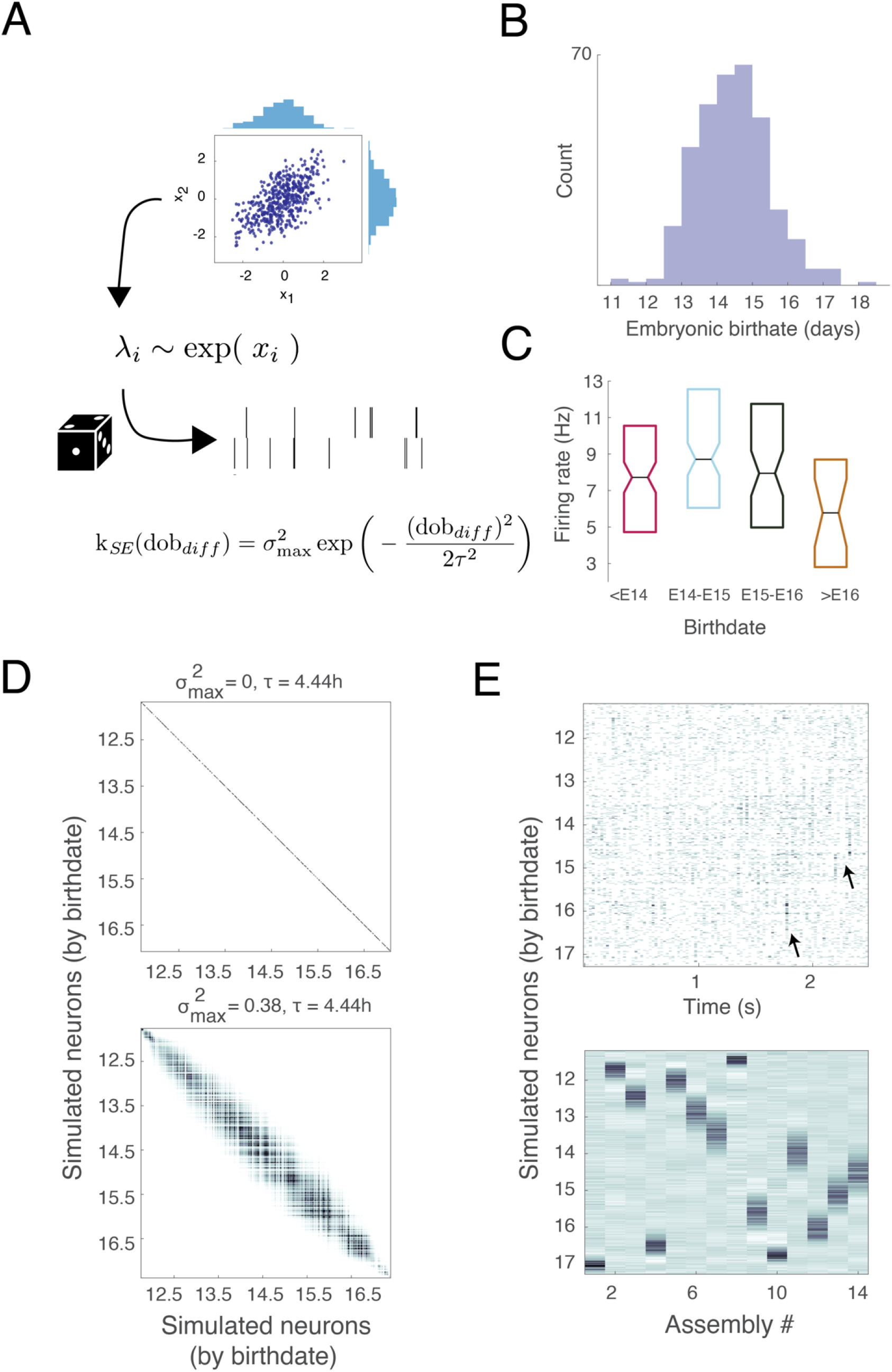
Linear-nonlinear Poisson model for exploring assembly dynamics. (**A**) Schematic illustrating the linear-nonlinear Poisson model. A multivariate Gaussian distribution x with predefined mean and covariance was transformed using an exponential nonlinearity. The resulting lognormally distributed process A was taken as the rate of a Poisson process to generate spike trains. *k*_*SE*_ (*dob*_*diff*_) is the kernel function for the Gaussian covariance, and depends on the difference of birthdates (*dob*_*diff*_) between simulated neurons. (**B**) Distribution of simulated birthdates of n=350 neurons. Birthdates were sampled from a Gaussian with mean birthdate (E)14.5, and standard deviation of 1 day. (**C**) Average firing rates of simulated neurons were set according to empirically observed firing rate distributions in SPW-Rs (**Fig. 3D**). (**D**) Analytical covariance matrices (*Cov*(λ_*i*_λ_*j*_)) for two values of 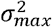, and a fixed value *τ*. These values were identical to the examples of simulated assembly expression rates in **Fig. 5C**. (**E**) Simulated raster plot (top) and extracted assembly independent components (bottom) with kernel parameters 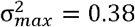= 0.38 and τ = 4.44*h*. Arrows in the raster plot point to observable assembly expressions.

**Supplementary Table 1.**
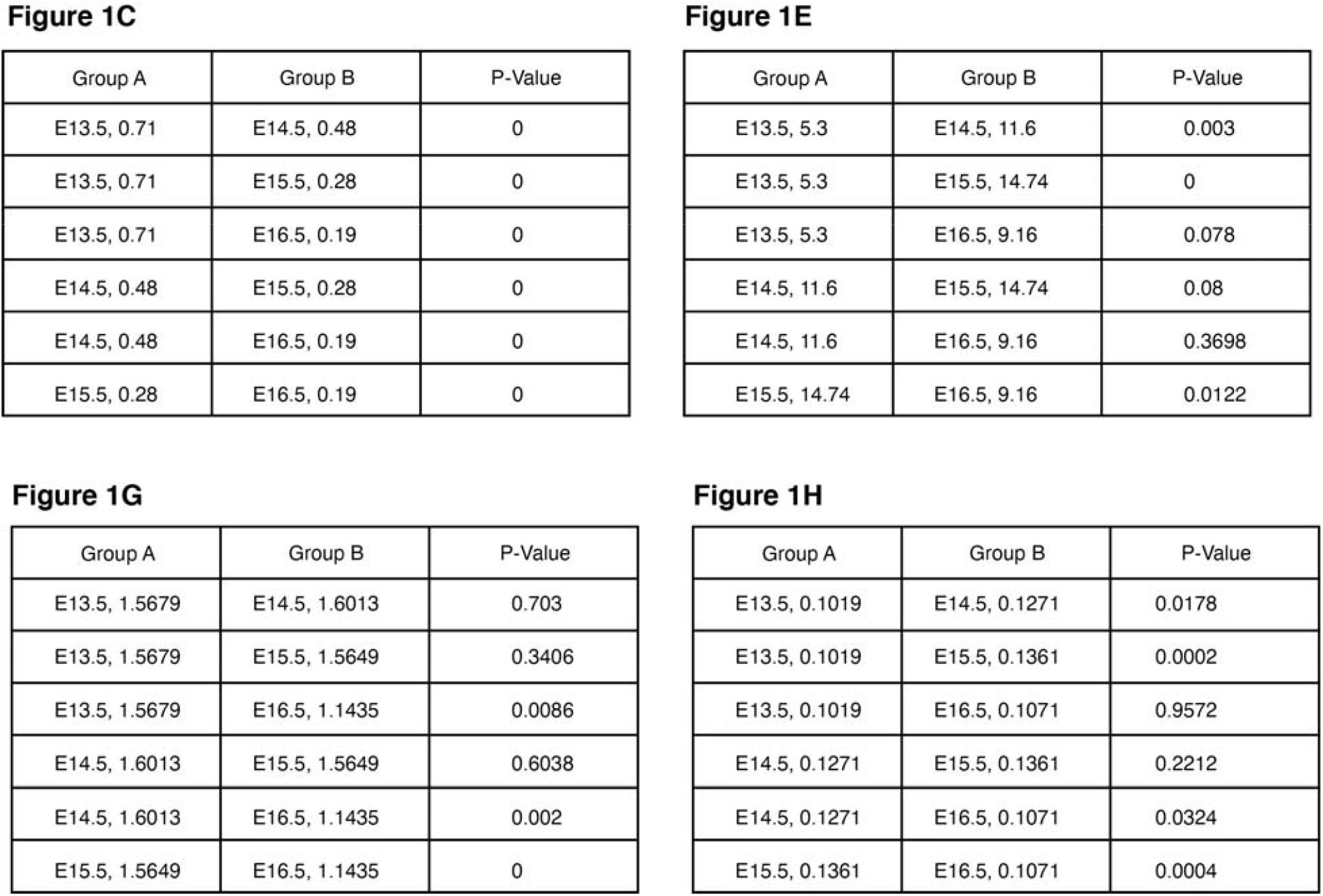
P-Values of multiple group comparisons pertaining to analyses of variance in Fig. 1. Values associated with each group are either means or medians, depending on the statistical test (see main figure legend).

**Figure 1C**
| Group A | Group B | P-Value |
| --- | --- | --- |
| E13.5, 0.71 | E14.5, 0.48 | 0 |
| E13.5, 0.71 | E15.5, 0.28 | 0 |
| E13.5, 0.71 | E16.5, 0.19 | 0 |
| E14.5, 0.48 | E15.5, 0.28 | 0 |
| E14.5, 0.48 | E16.5, 0.19 | 0 |
| E15.5, 0.28 | E16.5, 0.19 | 0 |

**Figure 1E**
| Group A | Group B | P-Value |
| --- | --- | --- |
| E13.5, 5.3 | E14.5, 11.6 | 0.003 |
| E13.5, 5.3 | E15.5, 14.74 | 0 |
| E13.5, 5.3 | E16.5, 9.16 | 0.078 |
| E14.5, 11.6 | E15.5, 14.74 | 0.08 |
| E14.5, 11.6 | E16.5, 9.16 | 0.3698 |
| E15.5, 14.74 | E16.5, 9.16 | 0.0122 |

**Figure 1G**
| Group A | Group B | P-Value |
| --- | --- | --- |
| E13.5, 1.5679 | E14.5, 1.6013 | 0.703 |
| E13.5, 1.5679 | E15.5, 1.5649 | 0.3406 |
| E13.5, 1.5679 | E16.5, 1.1435 | 0.0086 |
| E14.5, 1.6013 | E15.5, 1.5649 | 0.6038 |
| E14.5, 1.6013 | E16.5, 1.1435 | 0.002 |
| E15.5, 1.5649 | E16.5, 1.1435 | 0 |

| Group A | Group B | P-Value |
| --- | --- | --- |
| E13.5, 0.1019 | E14.5, 0.1271 | 0.0178 |
| E13.5, 0.1019 | E15.5, 0.1361 | 0.0002 |
| E13.5, 0.1019 | E16.5, 0.1071 | 0.9572 |
| E14.5, 0.1271 | E15.5, 0.1361 | 0.2212 |
| E14.5, 0.1271 | E16.5, 0.1071 | 0.0324 |
| E15.5, 0.1361 | E16.5, 0.1071 | 0.0004 |

**Supplementary Table 2.**
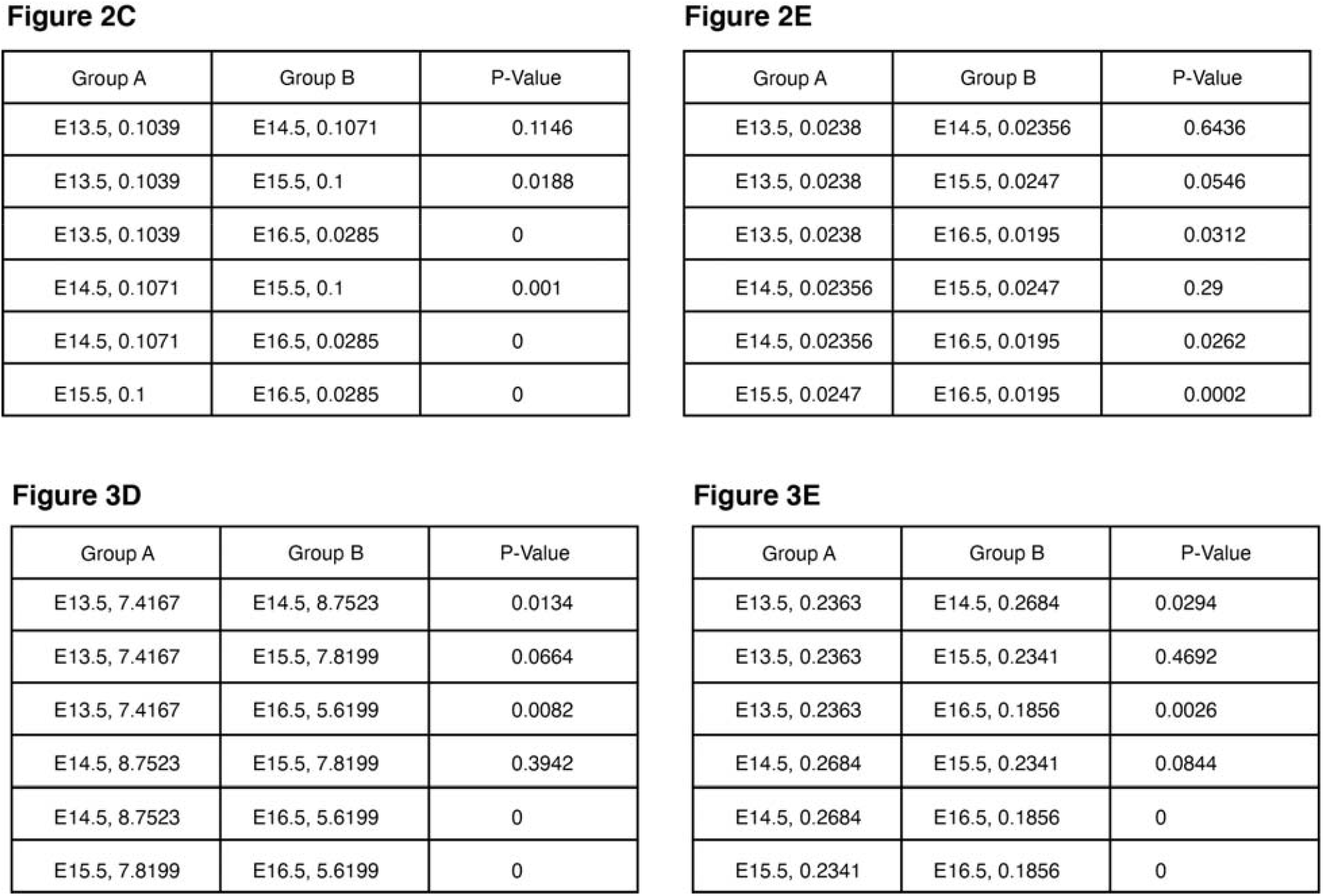
P-Values of multiple group comparisons pertaining to analyses of variance in **Fig. 2**,**3**. Values associated with each group are either means or medians, depending on the statistical test (see main figure legends).

**Figure 2C**
| Group A | Group B | P-Value |
| --- | --- | --- |
| E13.5, 0.1039 | E14.5, 0.1071 | 0.1146 |
| E13.5, 0.1039 | E15.5, 0.1 | 0.0188 |
| E13.5, 0.1039 | E16.5, 0.0285 | 0 |
| E14.5, 0.1071 | E15.5, 0.1 | 0.001 |
| E14.5, 0.1071 | E16.5, 0.0285 | 0 |
| E15.5, 0.1 | E16.5, 0.0285 | 0 |

**Figure 2E**
| Group A | Group B | P-Value |
| --- | --- | --- |
| E13.5, 0.0238 | E14.5, 0.02356 | 0.6436 |
| E13.5, 0.0238 | E15.5, 0.0247 | 0.0546 |
| E13.5, 0.0238 | E16.5, 0.0195 | 0.0312 |
| E14.5, 0.02356 | E15.5, 0.0247 | 0.29 |
| E14.5, 0.02356 | E16.5, 0.0195 | 0.0262 |
| E15.5, 0.0247 | E16.5, 0.0195 | 0.0002 |

**Figure 3D**
| Group A | Group B | P-Value |
| --- | --- | --- |
| E13.5, 7.4167 | E14.5, 8.7523 | 0.0134 |
| E13.5, 7.4167 | E15.5, 7.8199 | 0.0664 |
| E13.5, 7.4167 | E16.5, 5.6199 | 0.0082 |
| E14.5, 8.7523 | E15.5, 7.8199 | 0.3942 |
| E14.5, 8.7523 | E16.5, 5.6199 | 0 |
| E15.5, 7.8199 | E16.5, 5.6199 | 0 |

| Group A | Group B | P-Value |
| --- | --- | --- |
| E13.5, 0.2363 | E14.5, 0.2684 | 0.0294 |
| E13.5, 0.2363 | E15.5, 0.2341 | 0.4692 |
| E13.5, 0.2363 | E16.5, 0.1856 | 0.0026 |
| E14.5, 0.2684 | E15.5, 0.2341 | 0.0844 |
| E14.5, 0.2684 | E16.5, 0.1856 | 0 |
| E15.5, 0.2341 | E16.5, 0.1856 | 0 |

**Supplementary Table 3.**
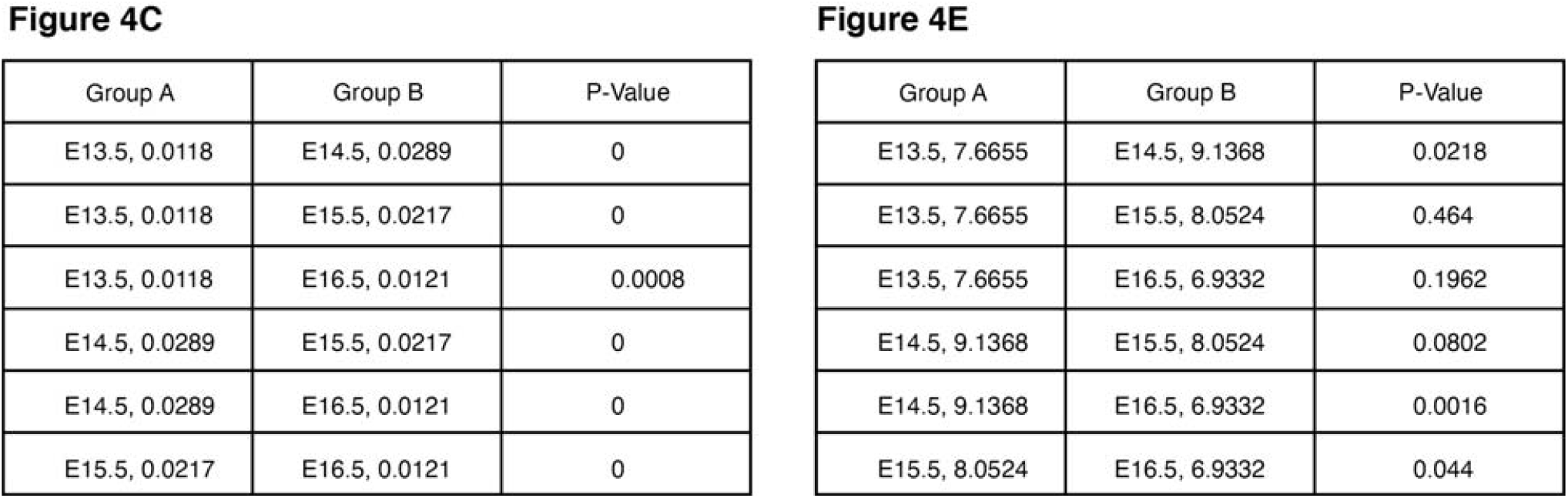
P-Values of multiple group comparisons pertaining to analyses of variance in **Fig. 4**. Values associated with each group are either means or medians, depending on the statistical test (see main figure legend).

**Figure 4C**
| Group A | Group B | P-Value |
| --- | --- | --- |
| E13.5, 0.0118 | E14.5, 0.0289 | 0 |
| E13.5, 0.0118 | E15.5, 0.0217 | 0 |
| E13.5, 0.0118 | E16.5, 0.0121 | 0.0008 |
| E14.5, 0.0289 | E15.5, 0.0217 | 0 |
| E14.5, 0.0289 | E16.5, 0.0121 | 0 |
| E15.5, 0.0217 | E16.5, 0.0121 | 0 |

| Group A | Group B | P-Value |
| --- | --- | --- |
| E13.5, 7.6655 | E14.5, 9.1368 | 0.0218 |
| E13.5, 7.6655 | E15.5, 8.0524 | 0.464 |
| E13.5, 7.6655 | E16.5, 6.9332 | 0.1962 |
| E14.5, 9.1368 | E15.5, 8.0524 | 0.0802 |
| E14.5, 9.1368 | E16.5, 6.9332 | 0.0016 |
| E15.5, 8.0524 | E16.5, 6.9332 | 0.044 |

**Supplementary Table 4.**
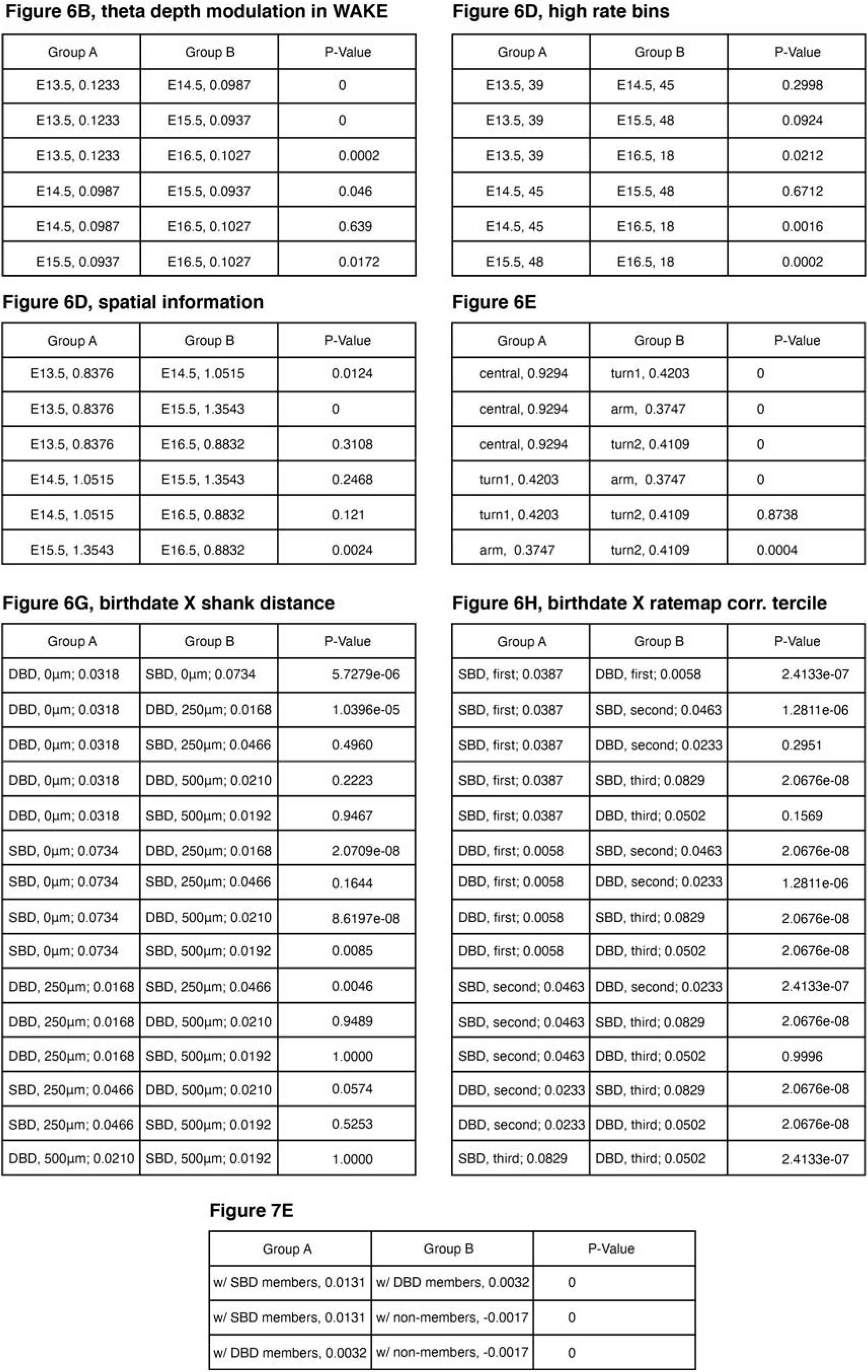
P-Values of multiple group comparisons pertaining to analyses of variance in **Fig. 6-7**. Values associated with each group are either means or medians, depending on the statistical test (see main figure legend).

**Figure 6B, theta depth modulation in WAKE**
| Group A | Group B | P-Value |
| --- | --- | --- |
| E13.5, 0.1233 | E14.5, 0.0987 | 0 |
| E13.5, 0.1233 | E15.5, 0.0937 | 0 |
| E13.5, 0.1233 | E16.5, 0.1027 | 0.0002 |
| E14.5, 0.0987 | E15.5, 0.0937 | 0.046 |
| E14.5, 0.0987 | E16.5, 0.1027 | 0.639 |
| E15.5, 0.0937 | E16.5, 0.1027 | 0.0172 |

**Figure 6D, high rate bins**
| Group A | Group B | P-Value |
| --- | --- | --- |
| E13.5, 39 | E14.5, 45 | 0.2998 |
| E13.5, 39 | E15.5, 48 | 0.0924 |
| E13.5, 39 | E16.5, 18 | 0.0212 |
| E14.5, 45 | E15.5, 48 | 0.6712 |
| E14.5, 45 | E16.5, 18 | 0.0016 |
| E15.5, 48 | E16.5, 18 | 0.0002 |

**Figure 6D, spatial information**
| Group A | Group B | P-Value |
| --- | --- | --- |
| E13.5, 0.8376 | E14.5, 1.0515 | 0.0124 |
| E13.5, 0.8376 | E15.5, 1.3543 | 0 |
| E13.5, 0.8376 | E16.5, 0.8832 | 0.3108 |
| E14.5, 1.0515 | E15.5, 1.3543 | 0.2468 |
| E14.5, 1.0515 | E16.5, 0.8832 | 0.121 |
| E15.5, 1.3543 | E16.5, 0.8832 | 0.0024 |

**Figure 6E**
| Group A | Group B | P-Value |
| --- | --- | --- |
| central, 0.9294 | turn1, 0.4203 | 0 |
| central, 0.9294 | arm, 0.3747 | 0 |
| central, 0.9294 | turn2, 0.4109 | 0 |
| turn1, 0.4203 | arm, 0.3747 | 0 |
| turn1, 0.4203 | turn2, 0.4109 | 0.8738 |
| arm, 0.3747 | turn2, 0.4109 | 0.0004 |

**Figure 6G, birthdate X shank distance**
| Group A | Group B | P-Value |
| --- | --- | --- |
| DBD, 0μm; 0.0318 | SBD, 0μm; 0.0734 | 5.7279e-06 |
| DBD, 0μm; 0.0318 | DBD, 250μm; 0.0168 | 1.0396e-05 |
| DBD, 0μm; 0.0318 | SBD, 250μm; 0.0466 | 0.4960 |
| DBD, 0μm; 0.0318 | DBD, 500μm; 0.0210 | 0.2223 |
| DBD, 0μm; 0.0318 | SBD, 500μm; 0.0192 | 0.9467 |
| SBD, 0μm; 0.0734 | DBD, 250μm; 0.0168 | 2.0709e-08 |
| SBD, 0μm; 0.0734 | SBD, 250μm; 0.0466 | 0.1644 |
| SBD, 0μm; 0.0734 | DBD, 500μm; 0.0210 | 8.6197e-08 |
| SBD, 0μm; 0.0734 | SBD, 500μm; 0.0192 | 0.0085 |
| DBD, 250μm; 0.0168 | SBD, 250μm; 0.0466 | 0.0046 |
| DBD, 250μm; 0.0168 | DBD, 500μm; 0.0210 | 0.9489 |
| DBD, 250μm; 0.0168 | SBD, 500μm; 0.0192 | 1.0000 |
| SBD, 250μm; 0.0466 | DBD, 500μm; 0.0210 | 0.0574 |
| SBD, 250μm; 0.0466 | SBD, 500μm; 0.0192 | 0.5253 |
| DBD, 500μm; 0.0210 | SBD, 500μm; 0.0192 | 1.0000 |

**Figure 6H, birthdate X ratemap corr. tercile**
| Group A | Group B | P-Value |
| --- | --- | --- |
| SBD, first; 0.0387 | DBD, first; 0.0058 | 2.4133e-07 |
| SBD, first; 0.0387 | SBD, second; 0.0463 | 1.2811e-06 |
| SBD, first; 0.0387 | DBD, second; 0.0233 | 0.2951 |
| SBD, first; 0.0387 | SBD, third; 0.0829 | 2.0676e-08 |
| SBD, first; 0.0387 | DBD, third; 0.0502 | 0.1569 |
| DBD, first; 0.0058 | SBD, second; 0.0463 | 2.0676e-08 |
| DBD, first; 0.0058 | DBD, second; 0.0233 | 1.2811e-06 |
| DBD, first; 0.0058 | SBD, third; 0.0829 | 2.0676e-08 |
| DBD, first; 0.0058 | DBD, third; 0.0502 | 2.0676e-08 |
| SBD, second; 0.0463 | DBD, second; 0.0233 | 2.4133e-07 |
| SBD, second; 0.0463 | SBD, third; 0.0829 | 2.0676e-08 |
| SBD, second; 0.0463 | DBD, third; 0.0502 | 0.9996 |
| DBD, second; 0.0233 | SBD, third; 0.0829 | 2.0676e-08 |
| DBD, second; 0.0233 | DBD, third; 0.0502 | 2.0676e-08 |
| SBD, third; 0.0829 | DBD, third; 0.0502 | 2.4133e-07 |

| Group A | Group B | P-Value |
| --- | --- | --- |
| w/ SBD members, 0.0131 | w/ DBD members, 0.0032 | 0 |
| w/ SBD members, 0.0131 | w/ non-members, -0.0017 | 0 |
| w/ DBD members, 0.0032 | w/ non-members, -0.0017 | 0 |

